# An Autoantigen Atlas from Human Lung HFL1 Cells Offers Clues to Neurological and Diverse Autoimmune Manifestations of COVID-19

**DOI:** 10.1101/2021.01.24.427965

**Authors:** Julia Y. Wang, Wei Zhang, Michael W. Roehrl, Victor B. Roehrl, Michael H. Roehrl

**Affiliations:** Curandis, New York, USA; Department of Gastroenterology, Affiliated Hospital of Guizhou Medical University, Guizhou, China; Department of Pathology and Human Oncology and Pathogenesis Program, Memorial Sloan Kettering Cancer Center, New York, USA; Human Oncology and Pathogenesis Program, Memorial Sloan Kettering Cancer Center, New York, USA

**Keywords:** COVID-19, SARS-Cov-2, autoantigens, autoantibodies, dermatan sulfate, autoimmunity

## Abstract

COVID-19 is accompanied by a myriad of both transient and long-lasting autoimmune responses. Dermatan sulfate (DS), a glycosaminoglycan crucial for wound healing, has unique affinity for autoantigens (autoAgs) from apoptotic cells. DS-autoAg complexes are capable of stimulating autoreactive B cells and autoantibody production. Using DS affinity, we identified an autoantigenome of 408 proteins from human fetal lung fibroblast HFL11 cells, at least 231 of which are known autoAgs. Comparing with available COVID data, 352 proteins of the autoantigenome have thus far been found to be altered at protein or RNA levels in SARS-Cov-2 infection, 210 of which are known autoAgs. The COVID-altered proteins are significantly associated with RNA metabolism, translation, vesicles and vesicle transport, cell death, supramolecular fibrils, cytoskeleton, extracellular matrix, and interleukin signaling. They offer clues to neurological problems, fibrosis, smooth muscle dysfunction, and thrombosis. In particular, 150 altered proteins are related to the nervous system, including axon, myelin sheath, neuron projection, neuronal cell body, and olfactory bulb. An association with the melanosome is also identified. The findings from our study illustrate a strong connection between viral infection and autoimmunity. The vast number of COVID-altered proteins with propensity to become autoAgs offers an explanation for the diverse autoimmune complications in COVID patients. The variety of autoAgs related to mRNA metabolism, translation, and vesicles raises concerns about potential adverse effects of mRNA vaccines. The COVID autoantigen atlas we are establishing provides a detailed molecular map for further investigation of autoimmune sequelae of the pandemic.

**Summary sentence:** An autoantigenome by dermatan sulfate affinity from human lung HFL1 cells may explain neurological and autoimmune manifestations of COVID-19

## Introduction

The emergence of the novel coronavirus SARS-CoV-2 has dragged the world into a prolonged pandemic. Aside from the intensively studied ACE2, heparan sulfate is another crucial entry receptor for coronaviruses (*1*). Dermatan sulfate (DS), structurally and functionally similar to heparan sulfate and heparin, belongs to the glycosaminoglycan family. Many viruses, including Ebola, Vaccinia, Zika, Dengue, and Hepatitis C viruses, have been shown to interact with glycosaminoglycans (*2–5*). These polyanionic polysaccharides consist of disaccharide repeating units of amino sugars and uronic acids with varying degrees of sulfation. Glycosaminoglycans are major components of the extracellular matrix and basement membrane, act as a filler between cells and tissue fibers and have numerous biological functions.

DS is most abundant in the skin but is also found in lungs, blood vessels, heart valves, and tendons. DS plays important roles in cell death, wound healing, and tissue repair. In human wound fluid, DS is the most abundant glycosaminoglycan (*6*). Its biosynthesis is increased by fibroblasts, epithelial cells, and capillary endothelial cells in wounded skin, mucosal ulcers, and inflammation-associated angiogenesis (*7–9*). Its molecular size also changes during wound healing, with elongated DS polymers packing along thin collagen fibrils in wounded skin (*10*). After tissue injury, fibroblasts require DS to migrate from the stroma surrounding the injury into the fibrin-laden wound to facilitate granulation tissue formation and wound healing (*11*).

DS is also a key molecule in autoimmunity, as we have discovered (*12–16*). DS is the most potent among glycosaminoglycans in stimulating autoreactive B1 cells and autoantibody production (*12, 13*). DS has a peculiar affinity to apoptotic cells and their released autoantigens (autoAgs), and macromolecular autoAg-DS affinity complexes are capable of engaging autoBCRs in a dual signaling event to activate B1 cells (*13, 14*). Recently, we also found that DS may steer autoreactive B1 cell fate at the pre-B stage by regulating the immunoglobulin heavy chain of the precursor BCR (*17*). Our studies illustrate a unifying property of autoAgs, i.e., self-molecules with DS affinity have a high propensity to become autoAgs, which explains how seemingly unrelated self-molecules can all induce humoral autoimmunity via similar immunological signaling events. In support of our hypothesis and by using DS affinity, we have cataloged hundreds of classic and novel autoAgs (*14–16, 18*).

A diverse spectrum of autoimmune symptoms has been observed in COVID-19 patients, including autoimmune cytopenia, multisystem inflammatory syndrome in children, immune-mediated neurological syndromes, Guillain-Barré syndrome, connective tissue disease-associated interstitial lung disease, antiphospholipid syndrome, autoimmune hemolytic anemia, autoimmune encephalitis, systemic lupus erythematosus, optic neuritis and myelitis, and acquired hemophilia (*19–26*). Many autoantibodies have been identified in COVID patients, including ANA (antinuclear antibody), ENA (extractable nuclear antigen), ANCA (anti-neutrophil cytoplasmic antibody), lupus anticoagulant, antiphospholipid, anti-IFN, anti-myelin oligodendrocyte glycoprotein, and anti-heparin-PF4 complex (*19–27*).

To understand autoimmune sequelae of COVID, we aimed to establish a COVID autoantigen atlas that will serve as a molecular map to guide further investigation. In this study, we identified an autoantigenome of 408 proteins from human fetal lung fibroblast HFL1 cells by DS-affinity fractionation and protein sequencing, with at least 231 being known autoAgs. We then compared these with currently available data from SARS-CoV-2-infected patients and cells (as of 12/14/2020 in Coronascape) (*28–48*). Remarkably, 352 (86.3%) of these proteins have been found to be altered (up- or down-regulated) at protein and/or RNA expression levels, and 210 of the COVID-altered proteins are known autoAgs in a great variety of autoimmune diseases and cancers. The COVID-altered proteins reveal intricate host responses to the viral infection and point to close associations with diverse disease manifestations of COVID-19.

## Results and Discussion

### An autoantigenome of 408 proteins with DS-affinity from HFL1 cells

Proteins extracted from HFL1 cells were fractionated with DS-affinity resins. The DS-binding fraction eluting with 0.5 M NaCl yielded 306 proteins by mass spectrometry sequencing, corresponding to proteins with medium-to-strong DS affinity. The fraction eluting with 1.0 M NaCl yielded 121 proteins, corresponding to proteins with very strong DS affinity. After excluding redundancies, a total of 408 unique proteins were obtained (Table 1). To verify how many of these proteins are known autoAgs, we conducted an extensive literature search for autoantibodies specific for each protein. Remarkably, at least 231 (56.6%) of our DS-affinity proteins already have known associated specific autoantibodies in various diseases and are thus confirmed autoAgs (see references in Table 1).

**Table 1.**
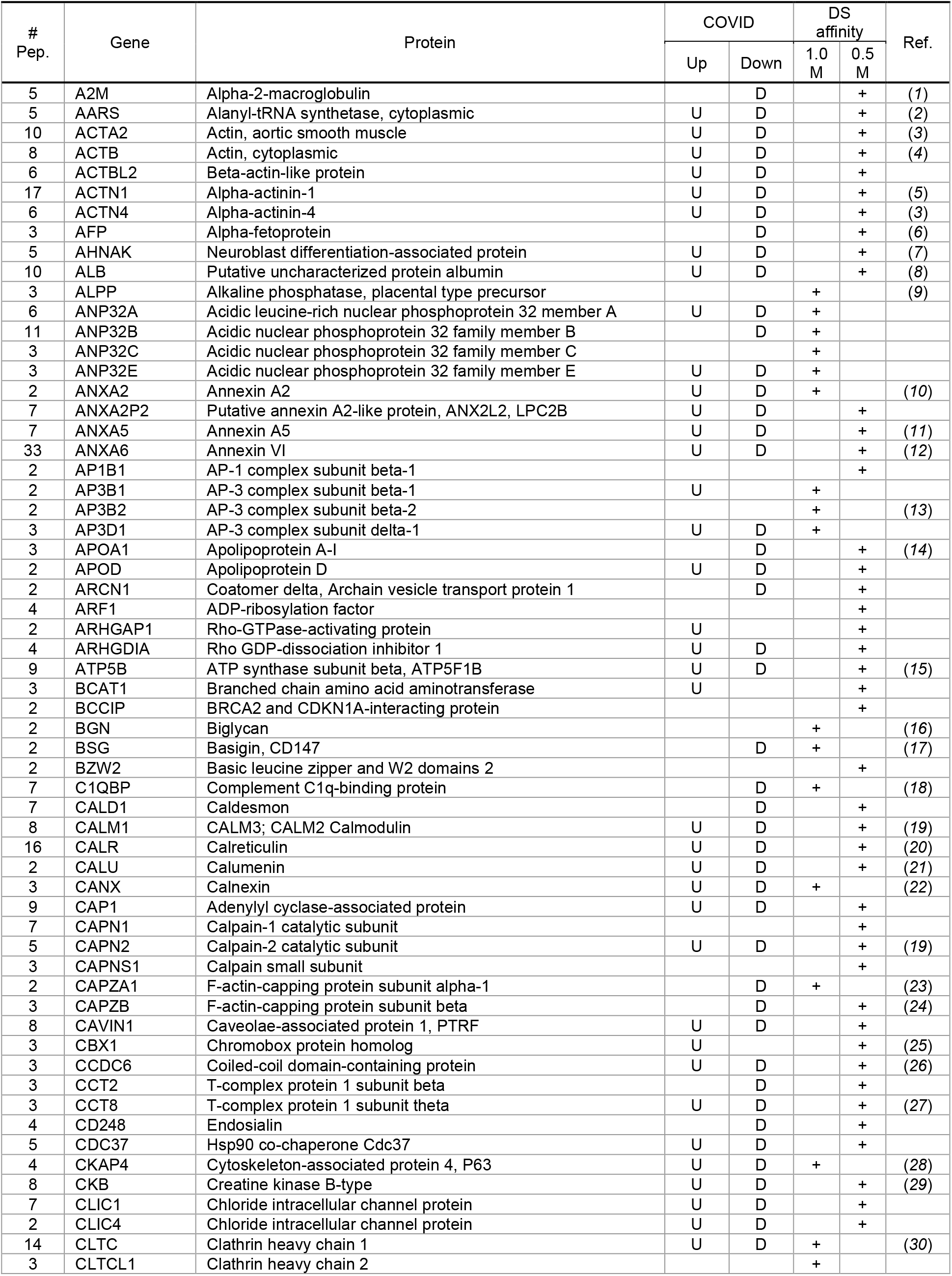
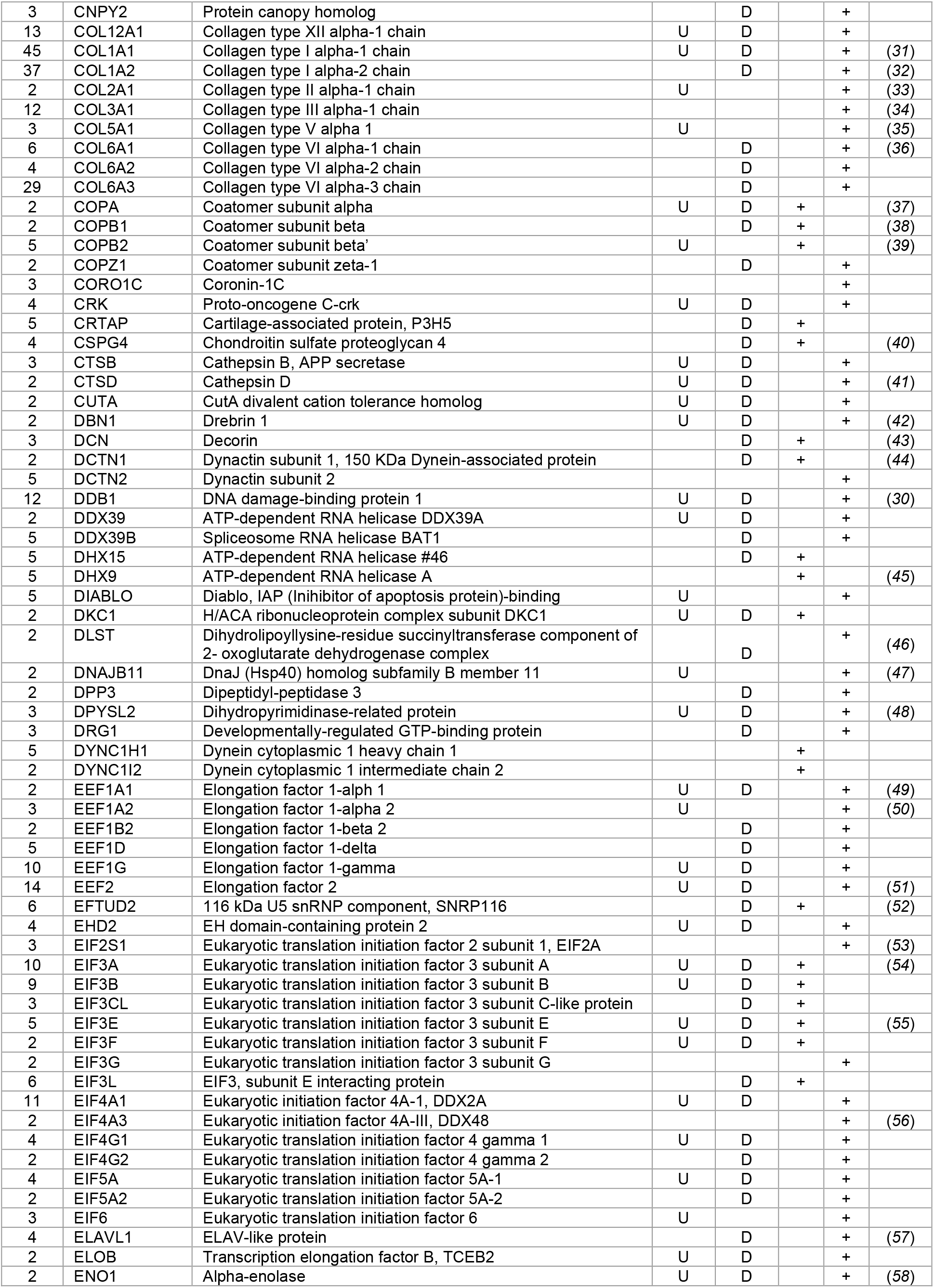
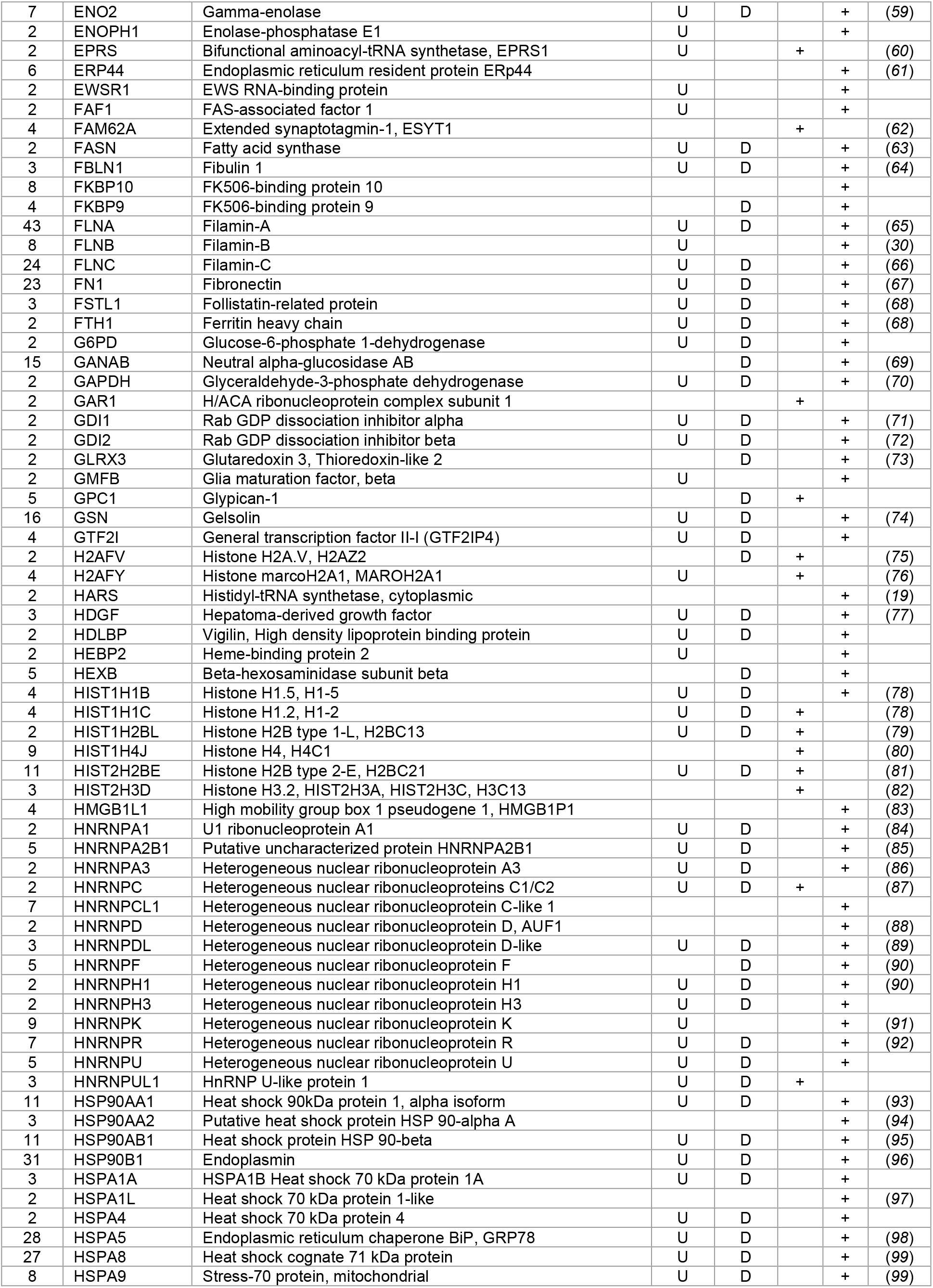
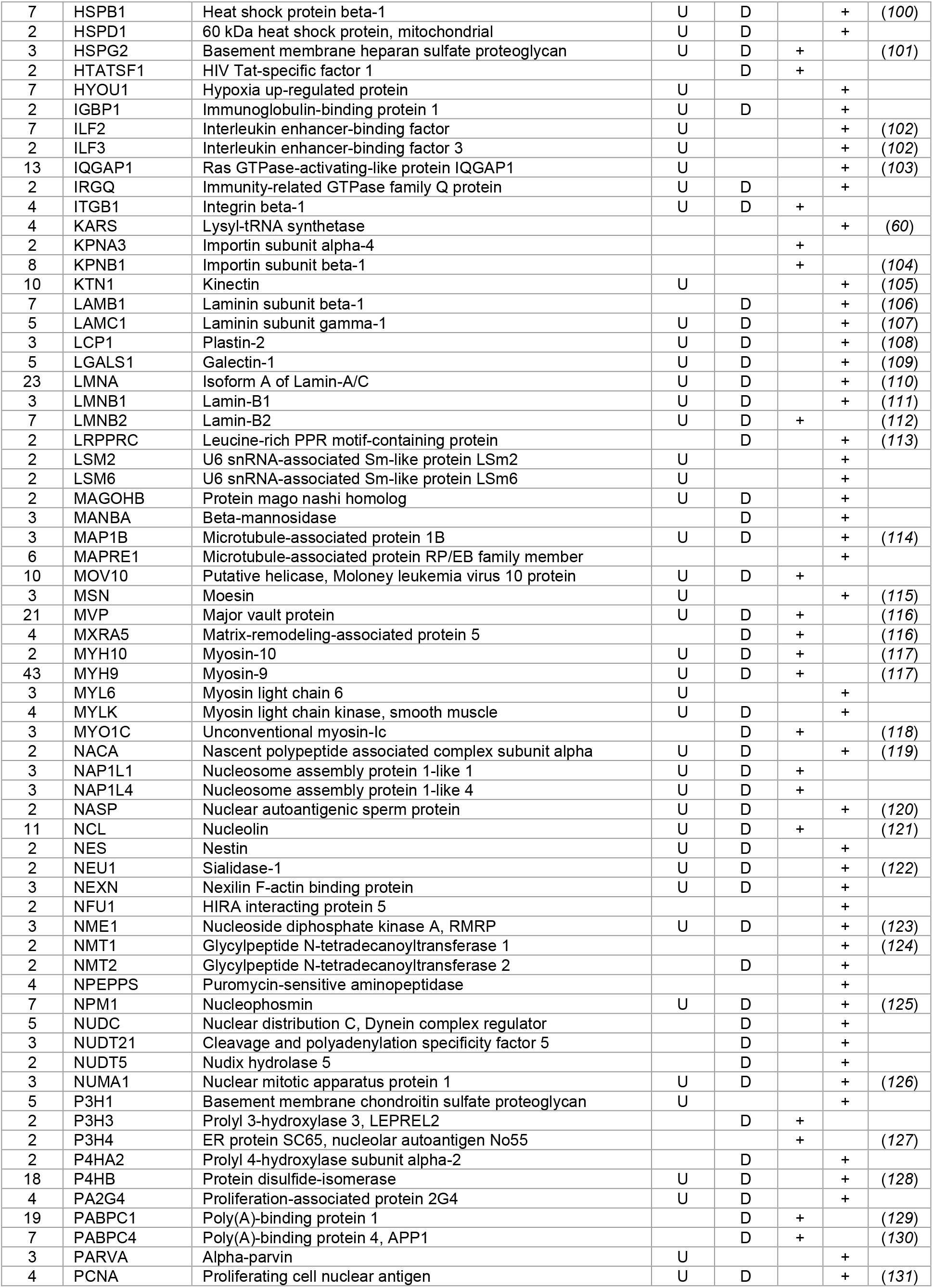
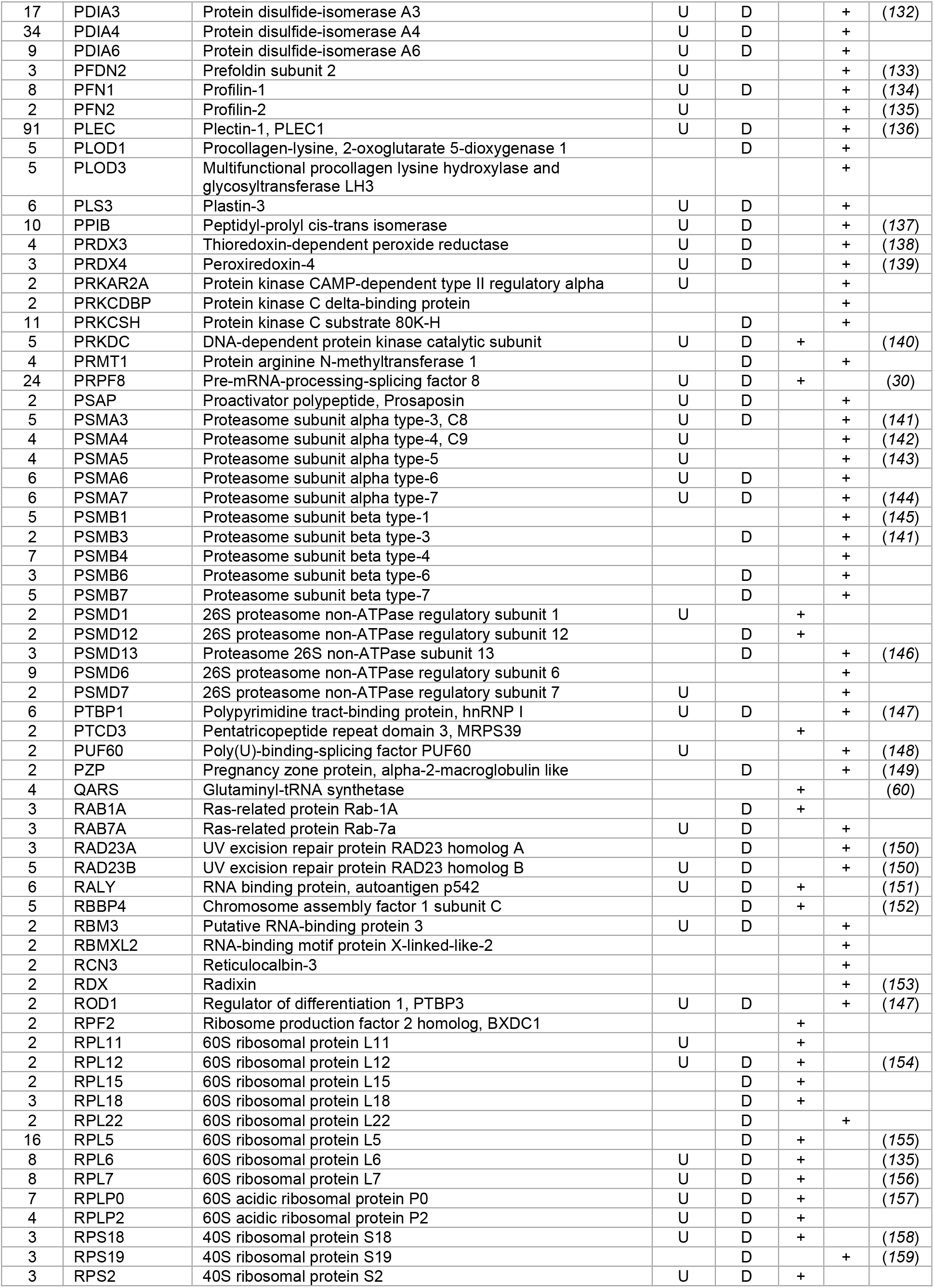
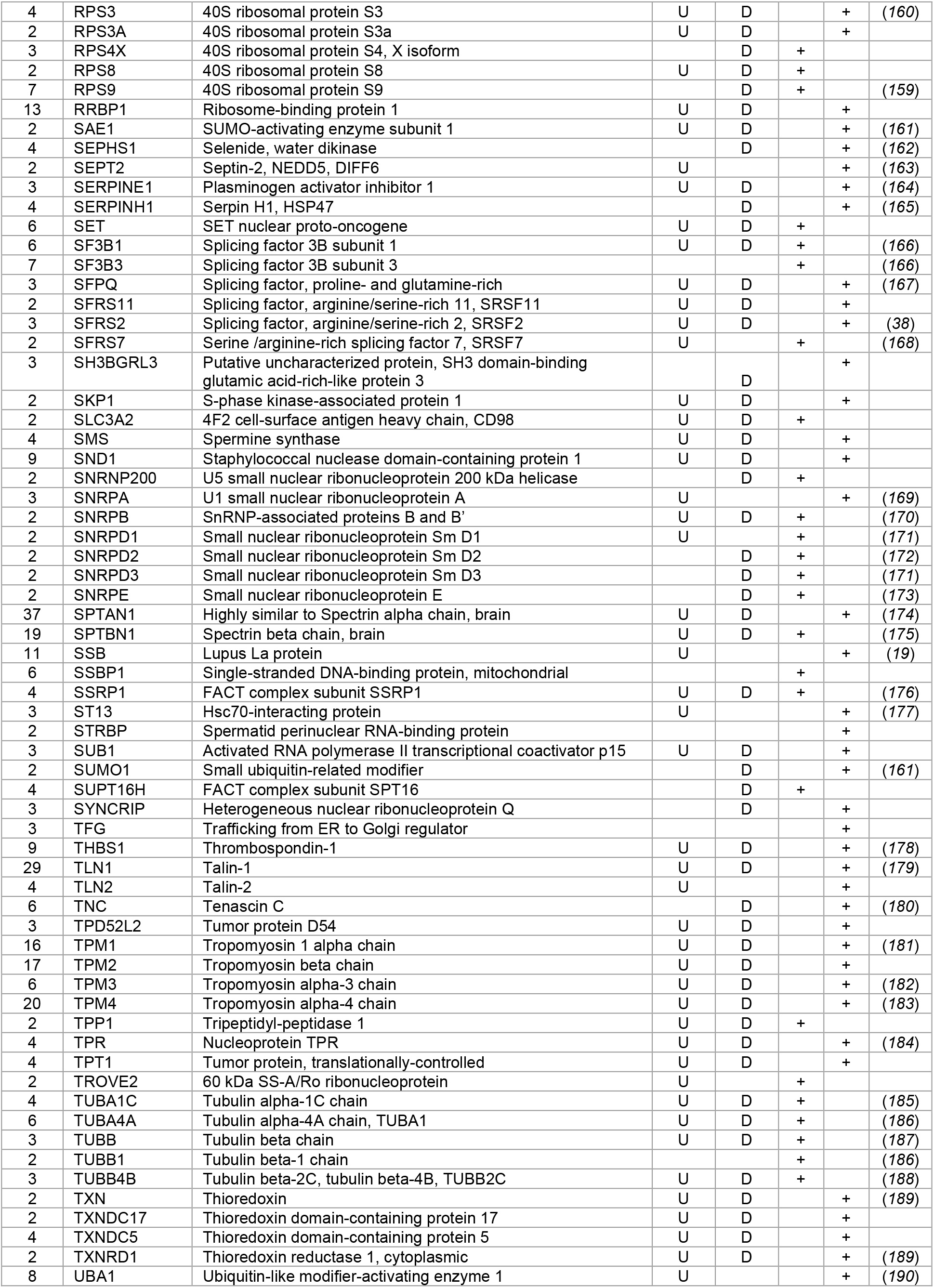
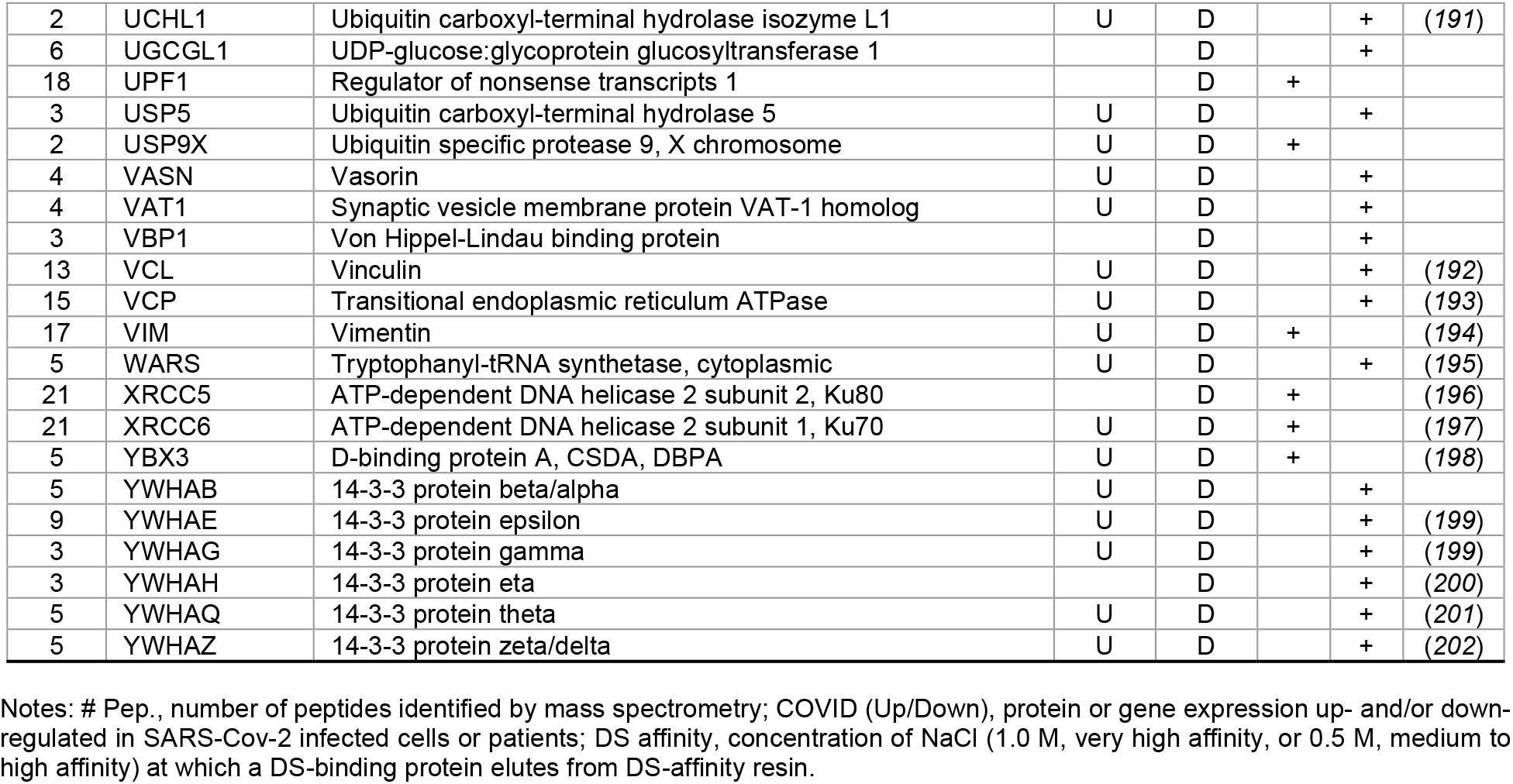
DS-affinity enriched autoantigenome from human HFL1 cells.

Of those not yet confirmed as autoAgs, a majority are similar to known autoAgs. As an example, we identified 18 ribosomal proteins, of which 9 have been individually identified as autoAgs (Table 1); however, anti-ribosomal autoantibodies are reported to react with a heterogeneous pool of many ribosomal proteins (*49*). Therefore, many of the ribosomal proteins we identified may be true but yet-to-be-confirmed autoAgs. As another example, autoantibodies against the 20S proteasome core are reported to be polyspecific and react with many subunits (*50*). Thus, although only 7 of 15 proteasome proteins we identified are thus far individually confirmed, the remainder may be true but yet-to-be-specified autoAgs. Similarly, some members of eukaryotic translation initiation and elongation factors are confirmed autoAgs, while others await confirmation. In summary, the putative autoantigenome from HFL1 cells provides at least 231 confirmed and 177 yet-to-confirm putative autoAgs (Table 1).

### DS-affinity proteins are functionally connected and enriched

To find out whether DS-affinity-associated proteins are a random collection or biologically connected, we performed protein-protein interaction analyses with STRING (*51*). Of the 408 DS-associated proteins, 405 proteins recognized by STRING (ANP32C, ANXA2P2, HSP90AA2 excluded) have 7,582 interactions, whereas a random set of 405 proteins is expected to have only 3,060 interactions; hence, DS-affinity proteins represent a significantly connected network with PPI enrichment p-value <1.0E-6 (Fig. 1). Based on cellular component classification, these proteins are highly concentrated in the nucleus (226 proteins), vesicles (111 proteins), ribonucleoprotein complexes (95 proteins), and the cytoskeleton (95 proteins).

**Fig. 1.**
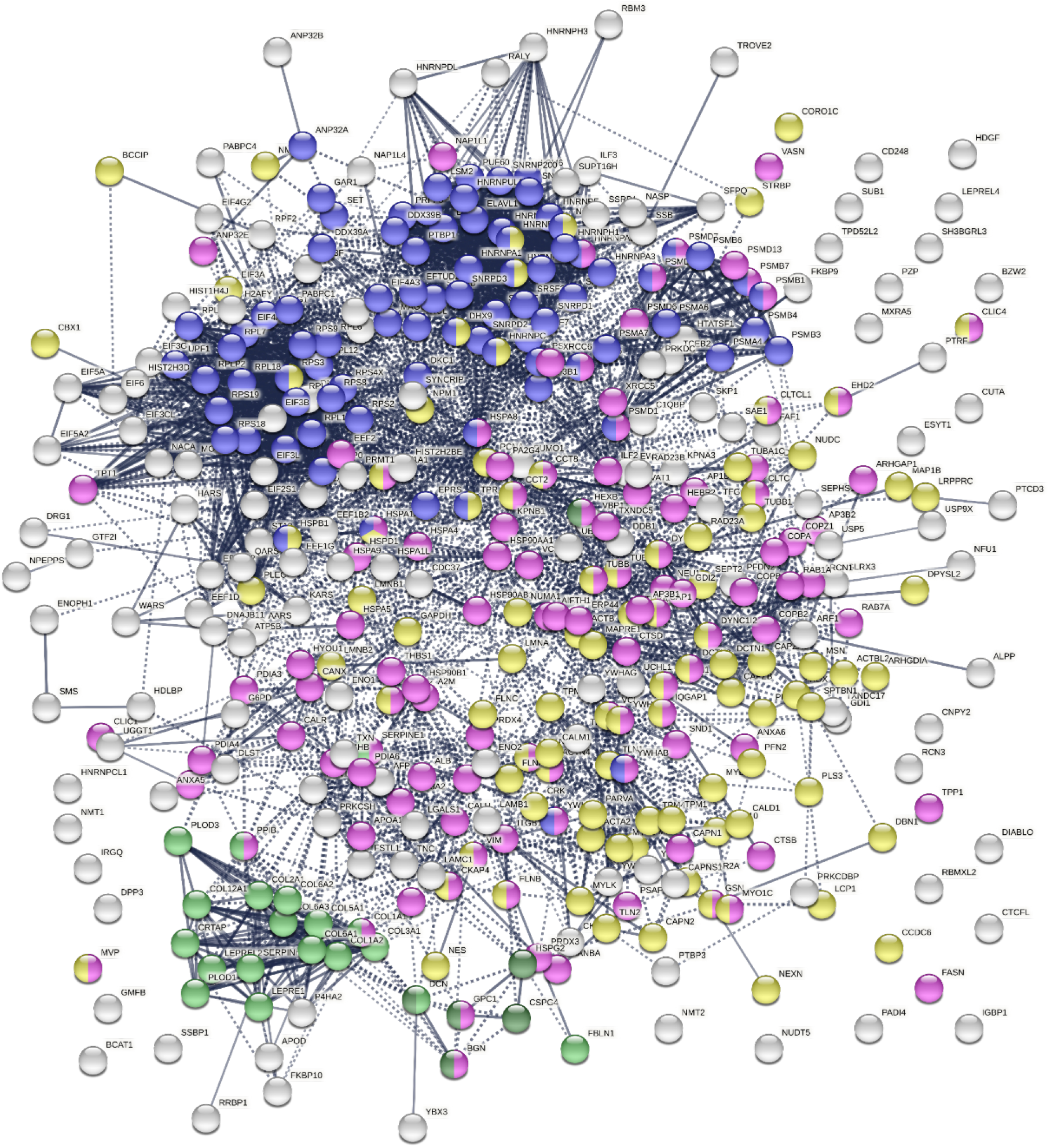
The 408-protein autoantigenome identified by DS-affinity from HFL1 cells forms a highly interacting network. Connecting lines represent interactions with high confidence (minimum interaction score of 0.7) as per STRING analysis. Colored proteins are involved in metabolism of RNA (blue), vesicles (pink), cytoskeleton (gold), collagen and elastic fibers (light green), and chondroitin sulfate/dermatan sulfate metabolism (darkgreen).

Pathway and process analyses by STRING and Metascape (*28*) revealed that the mRNA metabolic process is the most enriched GO Biological Process, and the top KEGG pathways are the spliceosome and protein processing in the endoplasmic reticulum. The top Reactome pathways are metabolism of RNA, metabolism of proteins, and axon guidance. The top local network clusters are GTP hydrolysis and joining of the 60S ribosomal subunits and mRNA splicing. The Molecular Complex Detection algorithm identified clusters related to eukaryotic translation elongation, cellular responses to stress, regulation of RNA stability, COPI-independent Golgi-to-ER retrograde traffic, and supramolecular fiber organization.

### 352 known and putative autoAgs are COVID-altered proteins

To find out which autoAgs may be involved in COVID-19, we compared the DS-affinity autoantigenome with proteins and genes that are up- or down-regulated in SARS-CoV-2 infection (Coronascape database comparison, Supplemental Table 1) (*28–48*). Remarkably, 352 (86.3%) of the 408 DS-affinity proteins have been found to be altered (up- and/or down-regulated at protein and/or mRNA levels) in COVID-19 patients or SARS-CoV-2 infected cells (Table 1). Of these, 260 are reported as up-regulated and 303 as down-regulated (including 211 that are both up- and down-regulated). The numbers are not conflicting, because the COVID data were generated by multiple proteomic and transcriptomic methods and different cells and tissues. A protein may not be overexpressed even when its mRNA is up-regulated, and a protein/gene may be up-regulated in one tissue or patient but down-regulated in another tissue or patient. A protein is considered altered if it is up- or down-regulated at the protein or RNA level and, in relation to SARS-CoV-2 infection, it is considered a COVID-altered protein.

Protein-interaction analysis revealed that 352 COVID-altered proteins form a highly connected network, exhibiting 6,286 interactions (vs. 2,451 expected; PPI enrichment p-value <1.0E-6) (Fig. 2). Based on cellular component analysis, the altered proteins can be located to intracellular organelles (323 proteins), nucleus (199 proteins), endomembrane system (143 proteins), vesicles (99 proteins), ribonucleoprotein complex (87 proteins), cytoskeleton (84 proteins), ER (72 proteins), and cell projections (52 proteins). Organelles with significant numbers of component proteins identified include the melanosome (30/105 proteins in melanosome), proteasome (16/64), polysome (13/66), spliceosome (34/187), ficolin-1-rich granule lumen (22/125), azurophil granules (17/155), and myelin sheath (26/157).

**Fig. 2.**
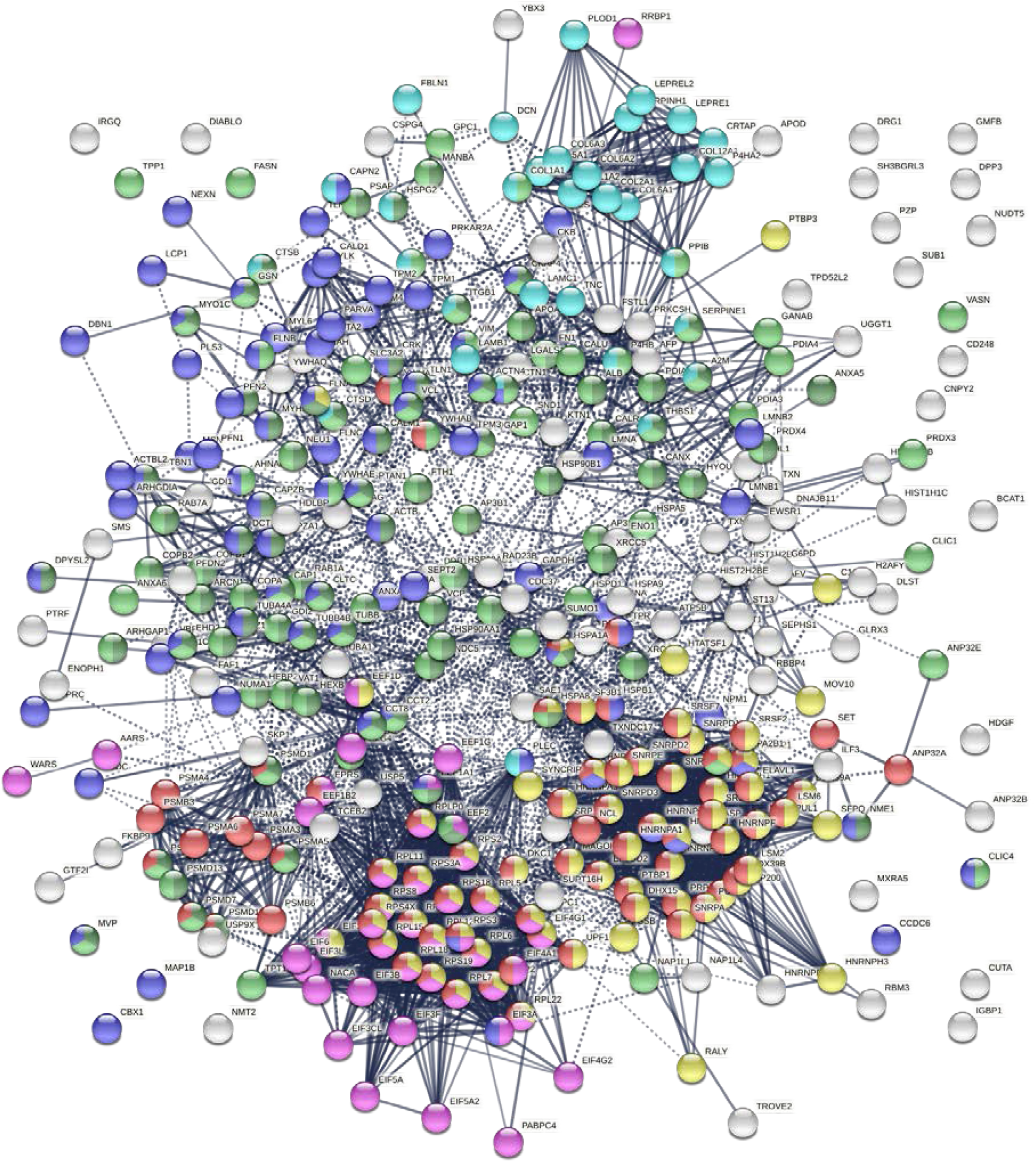
Network of 352 autoantigenome proteins that are altered in SARS-CoV-2 infected cells or patients. Connecting lines represent interactions with high confidence. Colored proteins are involved in metabolism of RNA (77 proteins, red), mRNA metabolic process (69 proteins, gold), translation (43 proteins, pink), vesicles (99 proteins, light green) and vesicle-mediated transport (84 proteins, dark green),cytoskeleton(84 proteins, blue),and extra cellular matrix organization(32 proteins, aqua).

Similarly, the group of 260 up-regulated proteins is highly connected (3,747 interactions vs. 1,424 expected) with significant enrichment in proteins associated with RNA and mRNA metabolism, translation, vesicles and vesicle-mediated transport, and regulation of cell death (Fig. 3A). The group of 303 down-regulated proteins is also highly connected (4,860 interactions vs. 1,907 expected), and these proteins are significantly related to RNA metabolism, translation, vesicles, cytoskeleton, and extracellular matrix (Fig. 3B).

**Fig. 3A.**
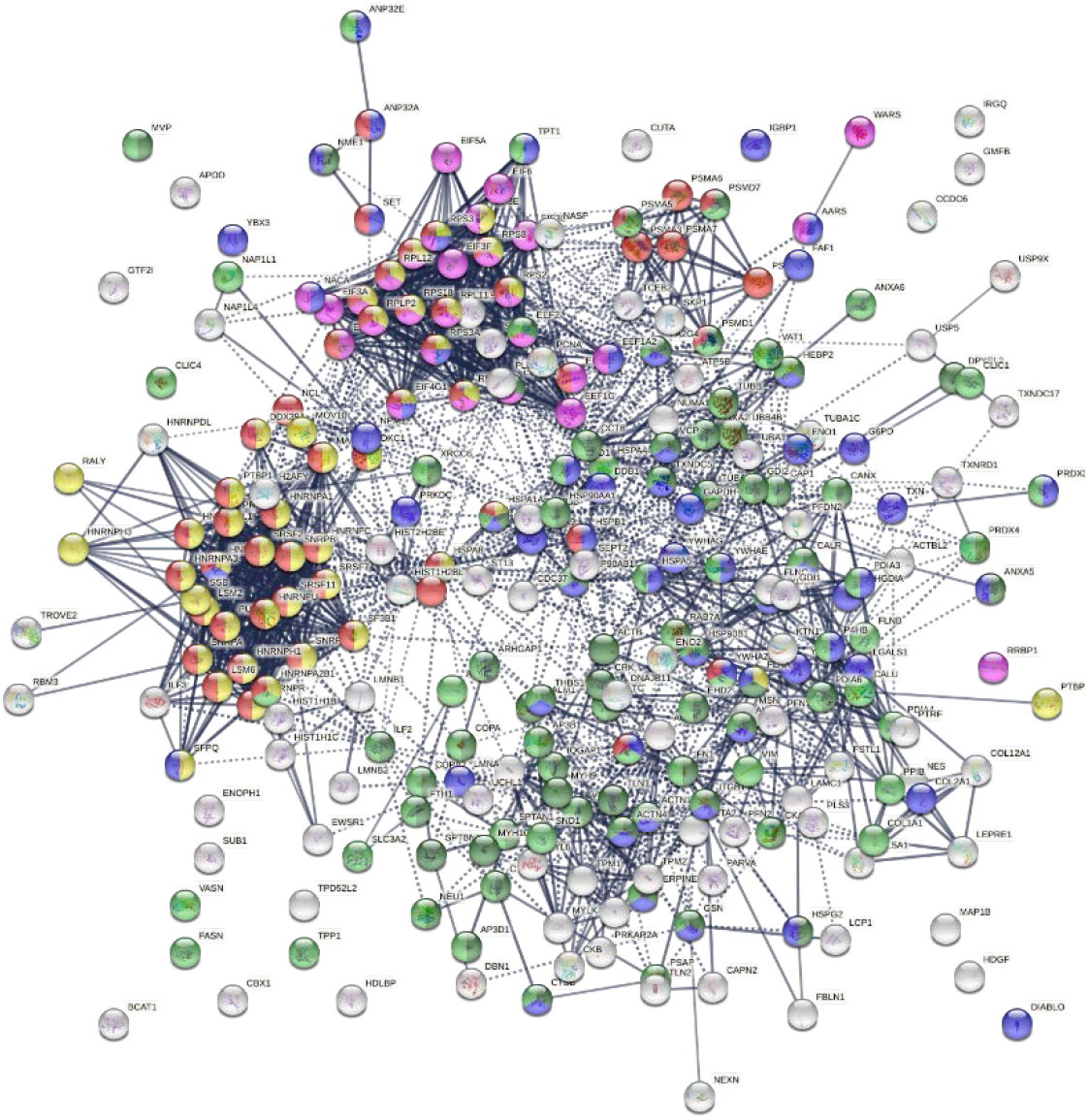
Interaction network of 260 up-regulated proteins in SARS-CoV-2 infected cells or patients. Connecting lines represent interactions with high confidence (minimum interaction score of 0.7). Colored proteins are involved in metabolism of RNA (54 proteins, red), translation (28 proteins, pink), vesicles (82 proteins, light green) and vesicle-mediated transport (67 proteins, dark green), regulation of cell death (61 proteins, blue), and mRNA metabolic process (46 proteins, gold).

**Fig. 3B.**
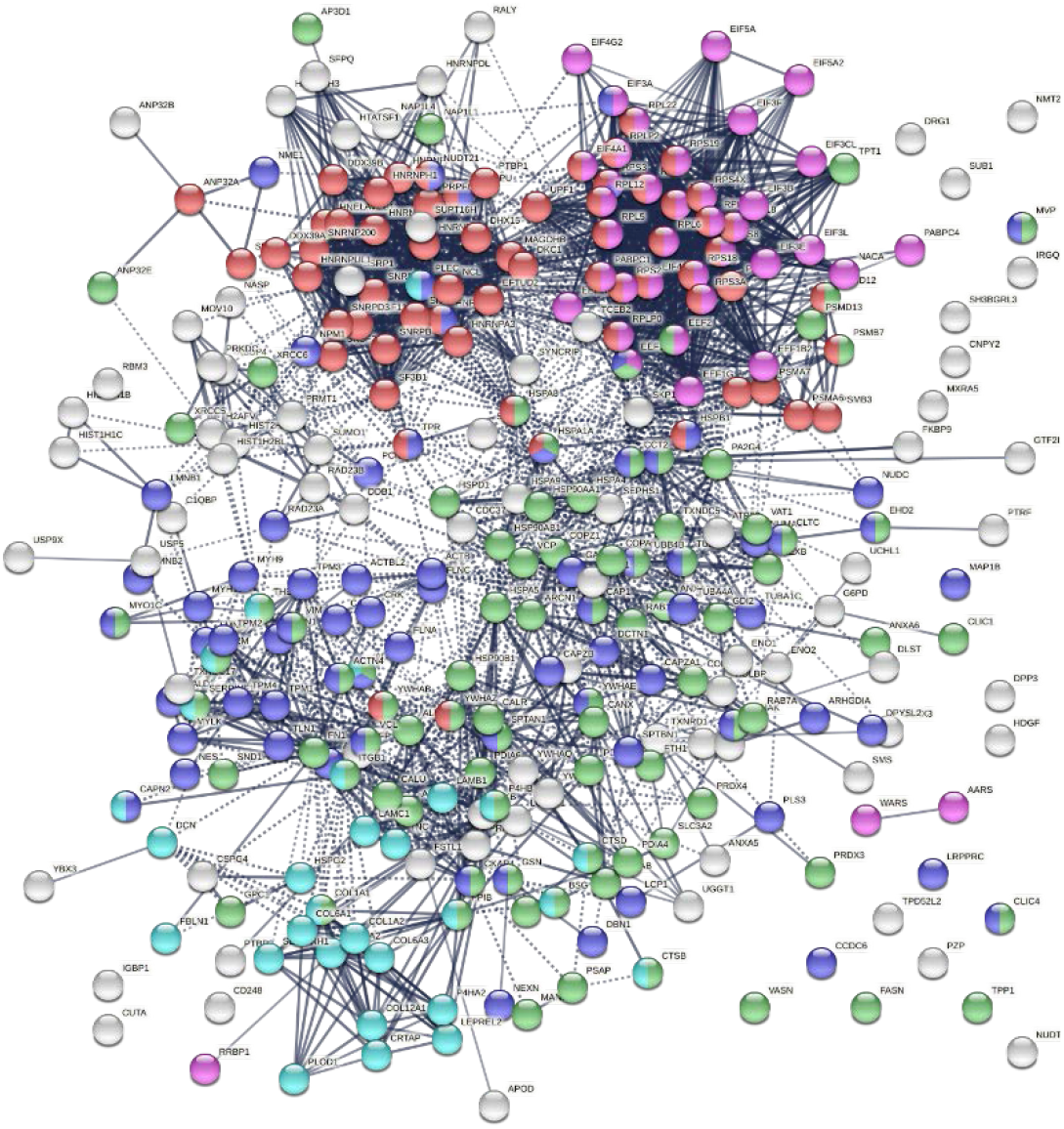
Interaction network of 303 down-regulated proteins in SARS-Cov-2 infected cells and patients. Connecting lines represent interactions with high confidence. Marked proteins are involved in RNA metabolism (64 proteins), translation (39 proteins, pink), vesicles (88 proteins, green), cytoskeleton (73 proteins, blue), and extracellular matrix organization (29 proteins, aqua).

### Pathways and processes affected by COVID-altered proteins

Network enrichment analysis by Metascape revealed that the 352 COVID-altered proteins are most significantly enriched in RNA metabolism, axon guidance, and translation (Table 2). Many processes, e.g., regulated exocytosis, wound healing, supramolecular fiber organization, smooth muscle contraction, and platelet degranulation are significantly affected by COVID-altered proteins regardless of whether they are up- or down-regulated. The up-regulated proteins are more related to axon guidance and interleukin signaling, whereas down-regulated proteins are more related to cellular response to stress and apoptosis.

**Table 2.** Top enriched pathways and processes related to COVID-altered proteins.

| COVID | Ontology | Description | Count | % | Log<sub>10</sub>(P) |
| --- | --- | --- | --- | --- | --- |
| **Altered** | R-HSA-8953854 | Metabolism of RNA | 78 | 22.16 | -51.2 |
|  | R-HSA-422475 | Axon guidance | 63 | 17.90 | -40.6 |
|  | GO:0006412 | Translation | 66 | 18.75 | -35.9 |
|  | GO:0000377 | RNA splicing | 44 | 12.50 | -28.0 |
|  | GO:0045055 | Regulated exocytosis | 58 | 16.48 | -26.7 |
|  | GO:0006457 | Protein folding | 33 | 9.38 | -24.3 |
|  | R-HSA-1474244 | Extracellular matrix organization | 33 | 9.38 | -20.6 |
|  | GO:0043687 | Post-translational protein modification | 35 | 9.94 | -20.0 |
|  | GO:0071826 | Ribonucleoprotein complex subunit organization | 32 | 9.09 | -19.7 |
|  | CORUM:5615 | Emerin complex 52 | 13 | 3.69 | -18.8 |
|  | GO:0010638 | Positive regulation of organelle organization | 40 | 11.36 | -16.1 |
|  | GO:0042060 | Wound healing | 38 | 10.80 | -15.6 |
|  | GO:0006913 | Nucleocytoplasmic transport | 30 | 8.52 | -15.6 |
|  | R-HSA-114608 | Platelet degranulation | 19 | 6.27 | -15.6 |
|  | R-HSA-5653656 | Vesicle-mediated transport | 40 | 11.36 | -15.4 |
|  | GO:0097435 | Supramolecular fiber organization | 41 | 11.65 | -15.1 |
|  | CORUM:1335 | SNW1 complex | 10 | 3.30 | -15.1 |
|  | GO:0002181 | Cytoplasmic translation | 18 | 5.11 | -15.1 |
|  | R-HSA-445355 | Smooth Muscle Contraction | 13 | 3.69 | -14.9 |
|  | GO:0031647 | Regulation of protein stability | 27 | 7.67 | -14.9 |
| **Up** | R-HSA-72163 | mRNA Splicing - Major Pathway | 23 | 8.85 | -18.8 |
|  | R-HSA-449147 | Signaling by Interleukins | 26 | 10.00 | -12.6 |
|  | GO:0000904 | Cell morphogenesis involved in differentiation | 31 | 11.92 | -11.3 |
| **Down** | R-HSA-2262752 | Cellular responses to stress | 55 | 18.15 | -34.5 |
|  | R-HSA-109581 | Apoptosis | 23 | 7.59 | -17.5 |
|  | GO:0035966 | Response to topologically incorrect protein | 22 | 7.26 | -15.0 |

Count: the number of DS-affinity proteins with membership in the given ontology term. %: percentage of DS-affinity proteins in the given ontology term.

### COVID-altered autoAgs are strongly related to the nervous system

COVID-19 patients frequently report neurological problems, such as loss of smell and taste, dizziness, headache, and stroke. While most symptoms are transient, some recovered patients are haunted by lingering neurological and psychological problems long after the viral infection. The underlying cause of transient and long-lasting neurological effects of COVID-19 has been puzzling. Analysis of COVID-altered proteins revealed a strong link to the nervous system. Of the 352 COVID-altered proteins, at least 150 are related to the nervous system (Fig. 4A). More than 60 proteins are related to axon guidance based on ontology analyses (Table 2 and Fig. 4A). In addition, there are 39 proteins related to neuron projection, 26 proteins related to myelin sheath, 25 proteins related to axon growth cone (*52*), 16 proteins related to neuronal cell body, 4 proteins related to cerebellar Purkinje cell layer, 3 proteins related to peripheral nervous system axon regeneration, and 2 proteins related to radial glial scaffolds. In particular, we found that 23 COVID-altered proteins are related to the olfactory bulb (*53*), which may explain the loss of smell in many COVID-19 patients.

**Fig. 4A.**
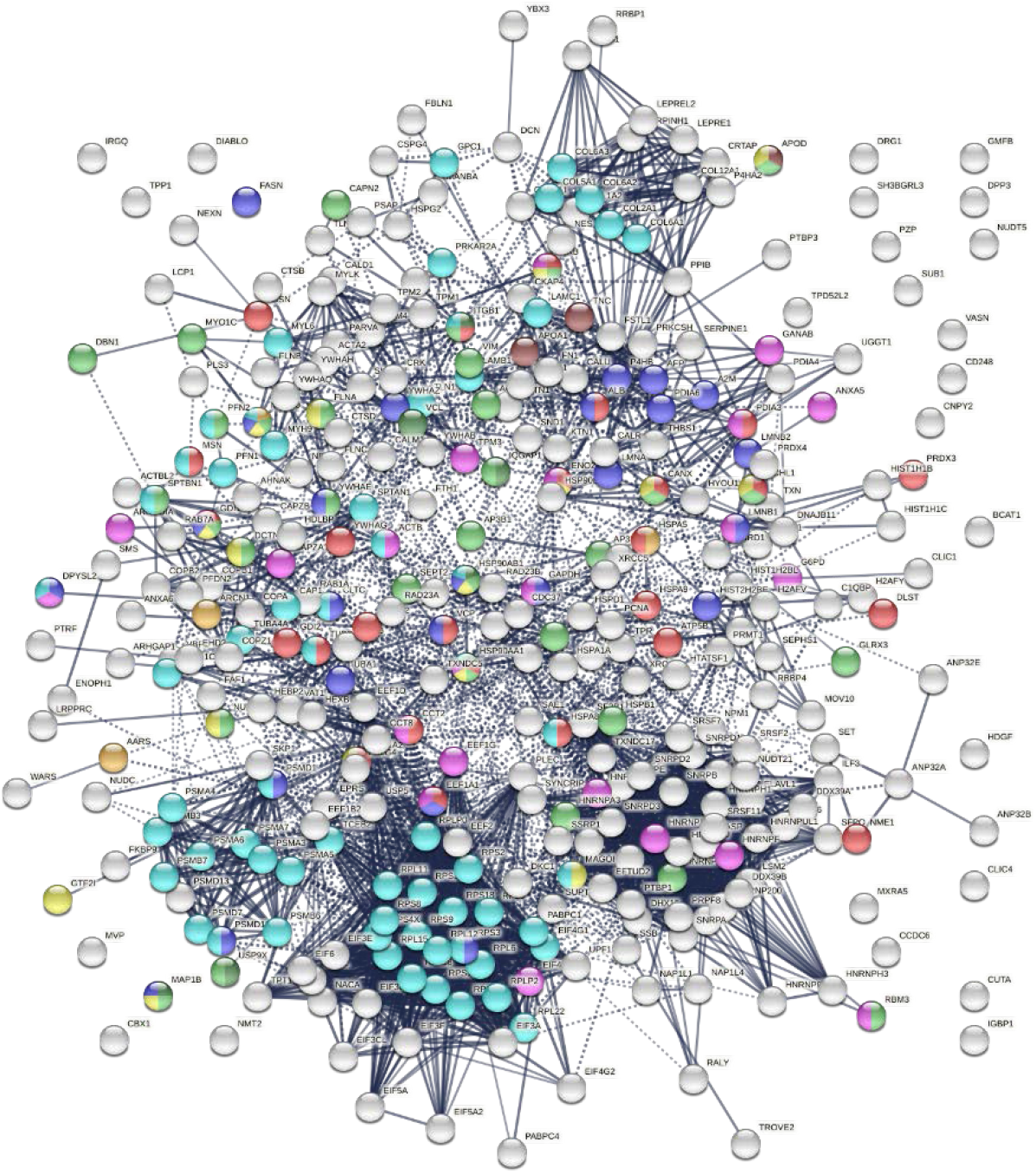
Nervous system-related proteins among COVID-altered proteins. Colored proteins are involved in axon guidance (62 proteins, aqua), axon growth cone (25 proteins, blue), myelin sheath (26 proteins, red), neuron projection (32 proteins, green) and neuron projection extension (7 proteins, dark green), neuronal cell body (16 proteins, gold), peripheral nervous system axon regeneration (3 proteins, brown), cerebellar Purkinje cell layer development (4 proteins, amber), and olfactory bulb (23 proteins, pink).

Most of these proteins are known autoAgs, e.g., ACTB, CANX, A2M, APOA1, CAPZA1, DPYSL2, FLNA, GDI2, LGALS1, MSN, PDIA3, PFN2, TNC, UCHL1, VCP, and VCL (see autoAg references in Table 1). Some yet-to-be-confirmed autoAgs with direct relation to the nerve system, e.g., NES (expressed mostly in nerve cells) and APOD (expressed by oligodendrocytes), warrant further investigation.

The COVID-altered proteins are also associated with a number of neurological diseases (Fig. 4B). By comparing our data with published proteomes, 23 proteins were similarly found in neuronal infection by Japanese encephalitis virus (*54*), 21 proteins in neuroblastoma (*55*), 22 proteins in glioblastoma (*56*), 26 proteins in neurodegeneration in Down syndrome (*57*), 22 proteins in Alzheimer disease hippocampus (*58*), 24 proteins in schizophrenia (*59*), 17 proteins in cerebral ischemia (*60*), and 17 proteins in Parkinson disease (*61*).

**Fig. 4B.**
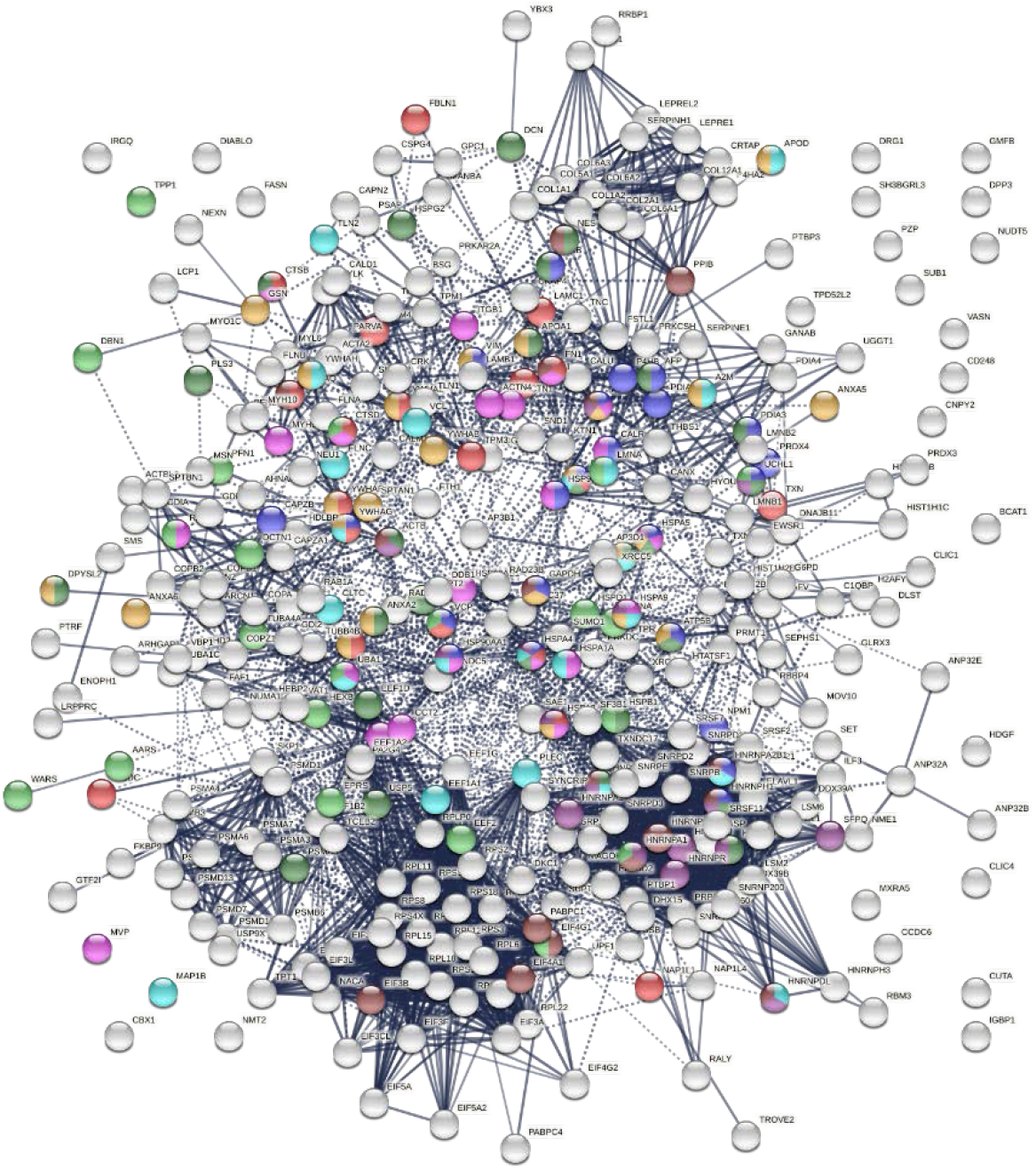
Neurological disease-related proteins among proteins altered in COVID. Colored are proteins found in neuronal infection with Japanese encephalitis virus (23 proteins, blue), neuroblastoma (21 proteins, red), glioblastoma (22 proteins, pink), neurodegeneration in Down syndrome (26 proteins, dark green), Alzheimer disease (22 proteins, aqua), schizophrenia (24 proteins, amber), cerebral ischemia induced neurodegenerative diseases (17 proteins, dark purple), Parkinson disease (17 proteins, brown),and neuro degeneration (21 proteins, green).

Coronavirus-induced demyelination has been reported in a mouse model of multiple sclerosis (*62*), which may explain our identification of 26 altered proteins related to the myelin sheath in SARS-CoV-2 infection. In a mouse brain injury model, DS appears to play an important role in glial scar formation and regeneration of dopaminergic axons (*63*). Alterations of white matter DS and extracellular matrix are specific, dynamic, and widespread in multiple sclerosis patients (*64*). DS has recently been reported to promote neuronal differentiation in mouse and human neuronal stem cells (*65*). Given the various functional roles of DS, our identification of a large number of known and putative autoAgs with DS affinity related to the nervous system is a compelling finding.

### COVID-altered autoAgs are related to cell death, wound healing, and blood coagulation

SARS-CoV-2 infection causes host cell death and leads to tissue injury. Wound healing, cellular response to stress, and apoptosis are among the most significant processes related to COVID-altered proteins (Table 2 and Fig. 5A). For example, we identified 66 proteins related to regulation of cell death and 23 related to regulation of apoptotic signaling pathways. DS binds to apoptotic cells and autoAgs released from dying cells, which has led to our previous identification of hundreds of autoAgs (*13–16, 18*). Upon tissue injury, DS biosynthesis is ramped up by fibroblasts and epithelial and endothelial cells (*7–9*). After tissue injury, DS assists fibroblast migration into the wound to facilitate granulation tissue formation and wound healing (*11*). DS, similar to heparin, is also an important anticoagulant that inhibits clot formation via interaction with antithrombin and heparin cofactor II (*66*). Given these biological roles of DS, it is consistent that a large number of COVID-altered proteins related to cell death and tissue injury are identified by DS-affinity.

**Fig. 5A.**
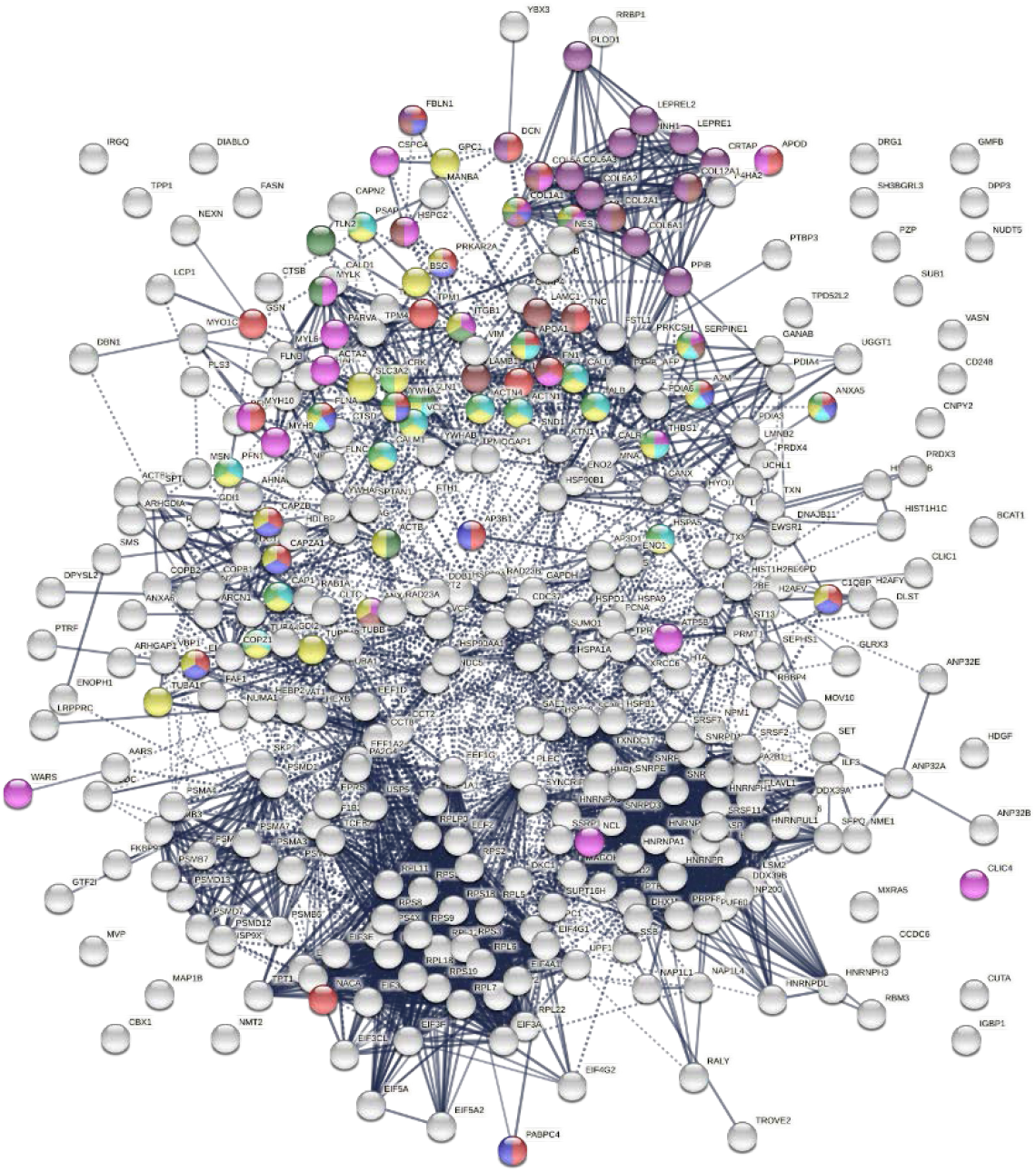
Relation of COVID-altered proteins to wound healing and hemostasis. Response to wounding (25 proteins, red), blood vessel development (20 proteins, pink), blood coagulation (14 proteins, blue), collagen-containing extracellular matrix (13 proteins, brown), collagen biosynthesis and modifying enzymes (16 proteins, dark purple), platelet activation (3 proteins, dark green) and platelet activation signaling and aggregation (22 proteins, green), platelet degranulation (18 proteins, aqua), and hemostasis (35 proteins, gold).

Blood coagulation and thrombosis are frequent complications of COVID-19. Platelet degranulation is found to be significantly associated with at least 18 altered proteins (Table 2 and Fig. 5A). COVID-altered proteins are related to blood coagulation, platelet activation, platelet alpha granules, fibrinogen binding, fibrinogen complex, platelet plug formation, von Willebrand factor A-like domain superfamily, and platelet-derived growth factor binding. Collagens, which support platelet adhesion and activation, and collagen biosynthesis and modifying enzymes are also among the COVID-altered proteins, e.g., collagen type VI trimer and type I trimer (Fig. 5A). The majority of these altered proteins are known autoAgs, e.g., ALB, ANXA5, C1QBP, CALM1, CAPZB, COL1A1, COL1A2, COL6A1, FBLN1, FN1, PLEC, PPIB, THBS1, TLN1, TUBA4A, and YWHAZ (see autoAg references in Table 1). Some are unknown and await further investigation, e.g., AP3B1, CRK, CTSB, EHD2, PLOD1, PSAP, and PARKAR2A.

### Supramolecular fibril alteration offers clues to muscle dysfunction and fibrosis

Over 50 supramolecular filament proteins are identified by DS-affinity from HFL1 cells. Remarkably, nearly all (except for one) are found to be altered in SARS-CoV-2 infection, and the majority have already been reported as autoAgs (Table 1). They include various isoforms of actin, actinin, collagen, filamin, fibronectin, fibulin, dynactin, dynein, lamin, myosin, nestin, nexilin, profilin, plectin, plastin, proteoglycan, septin, spectrin, talin, tropomyosin, tubulin, vinculin, and vimentin (Table 1, Fig. 5B). These proteins are major components of the extracellular matrix, basement membrane, cell cytoskeleton, cytoskeletal motors, muscle filaments, and contractile motors of muscle cells.

**Fig. 5B.**
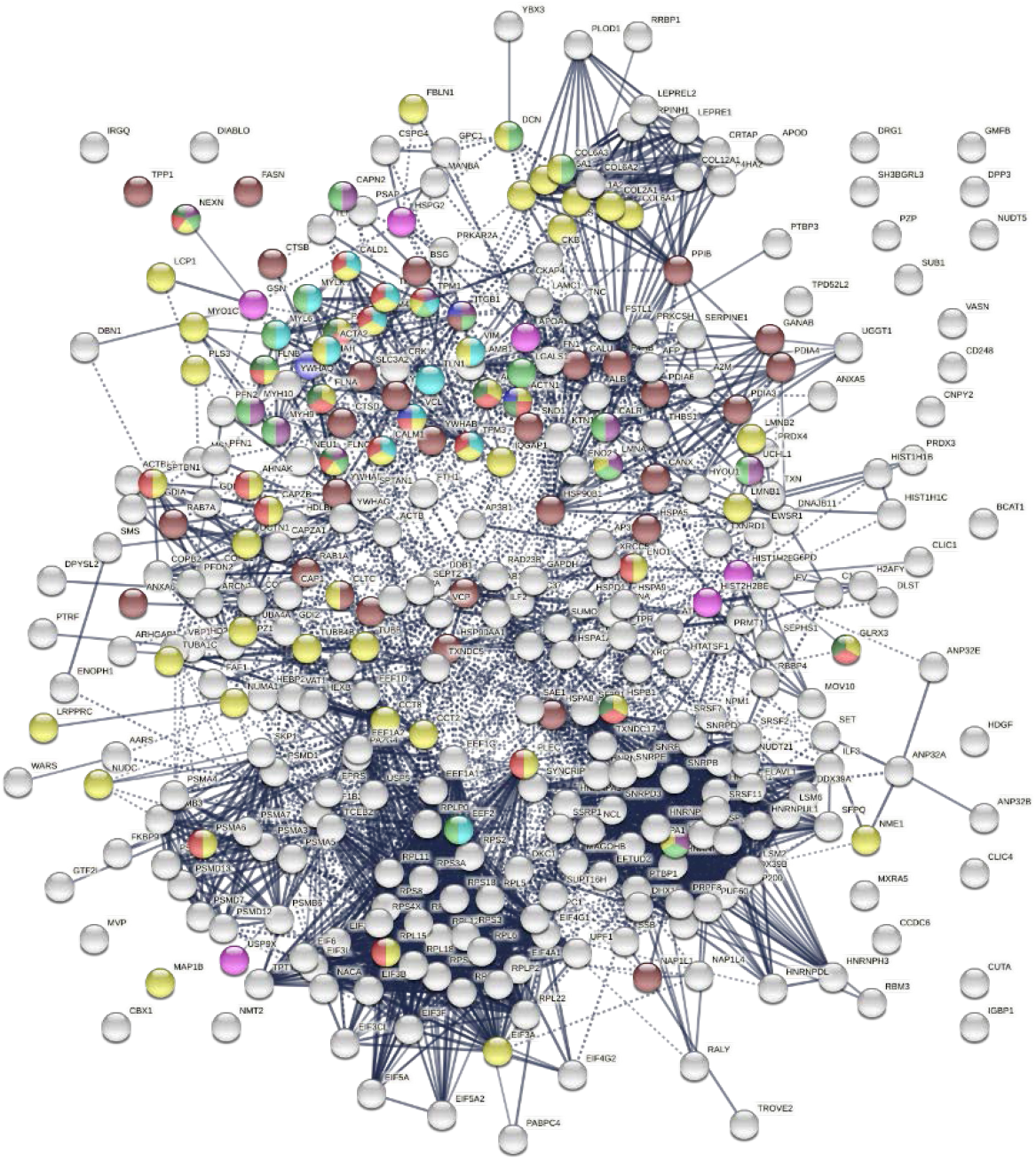
Other significantly enriched groups among altered proteins. Supramolecular fiber (56 proteins, amber), melanosome (30 proteins, brown), striated muscle cell differentiation (11 proteins, purple), myofibril (23 proteins, red), muscle structure development (18 proteins, green), muscle contraction (13 proteins, aqua), Z disk (9 proteins, dark green), intercalated disk (4 proteins, blue), and amyloid fiber formation(6 proteins, pink).

A significant number of COVID-altered proteins are related. Emerin complex and smooth muscle contraction are among the top enriched biological processes of COVID-altered proteins (Table 2, Fig. 5B). Emerin is highly expressed in cardiac and skeletal muscle, and emerin mutations cause X-linked recessive Emery-Dreifuss muscular dystrophy, cardiac conduction abnormalities, and dilated cardiomyopathy. Smooth muscle resides primarily in the walls of hollow organs where it performs involuntary movements, e.g., respiratory tract, blood vessels, gastrointestinal tract, and renal glomeruli. In addition, we identified proteins with significant association to myofibrils (the contractile elements of skeletal and cardiac muscle; 23 proteins) (Fig. 5B), stress fiber (a contractile actin filament bundle that consists of short actin filaments with alternating polarity: MYH9, MYLK, FLNB, TPM1, TPM2, TPM3, TPM4, ACTN1, ACTN4), muscle filament sliding (the sliding of actin thick filaments and myosin thick filaments past each other in muscle contraction), Z disk (plate-like region of a muscle sarcomere to which the plus ends of actin filaments are attached), intercalated disc (a cell-cell junction complex at which myofibrils terminate in cardiomyocytes, mediates mechanical and electrochemical integration between individual cardiomyocytes), and negative regulation of smooth muscle cell-matrix adhesion (2 proteins; SERPINE1, APOD).

Pulmonary fibrosis is prominent in COVID-19 and contributes to lethality in some cases (*67, 68*). Fibrosis, or fibrotic scarring, is pathological wound healing in which excessive extracellular matrix components are produced by fibroblasts and accumulate in the wounded area. Histopathological examination of COVID-19 patients found highly heterogenous injury patterns reminiscent of exacerbation of interstitial lung disease, including interstitial thickening, fibroblast activation, and deposition of collagen fibrils (*22*). We identified a significant number of COVID-altered proteins that are associated with collagen bundles and collagen biosynthesis and modifying enzymes (16 proteins), extracellular matrix organization (33 proteins), supramolecular fibers, and amyloid formation offering functional links to fibrosis (Fig. 5B).

### Potential autoAgs in COVID-19 patients and a connection to the melanosome

To find out how altered proteins may differ in patients, we compared our putative autoantigenome to published single-cell RNA sequencing data of 6 patients hospitalized for COVID-19 (*28, 34*) and identified 32-59 putative autoAgs per patient (Fig. 6). Interestingly, while identified from different patients, the altered proteins/genes identified share involvement of leukocyte activation, vesicles and vesicle transport, protein processing in the ER (including antigen processing and presentation), regulation of cell death, translation, muscle contraction, myelin sheath, and curiously, the melanosome (Fig. 6). The estrogen signaling pathway and thyroid hormone synthesis are found to be associated with altered proteins in some patients. Patient C2 has 5 altered proteins related to neuron differentiation regulation, and patient C4 has 6 altered proteins related to neuron death.

**Fig. 6.**
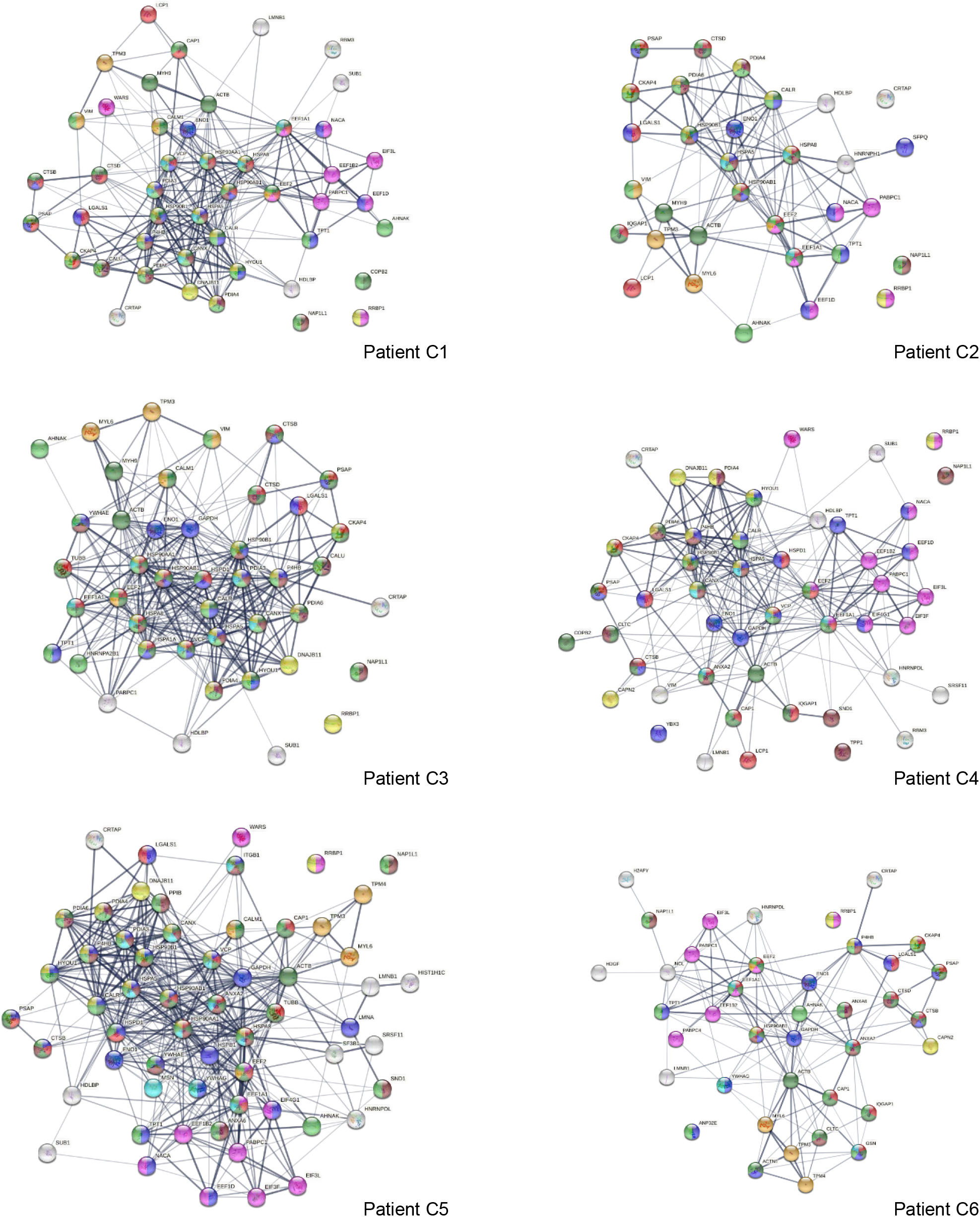
Interaction network of altered proteins in 6 COVID-19 patients. Colored proteins are associated with leukocyte activation involved in immune response (red), vesicles (light green) and vesicle-mediated transport (dark green), protein processing in the ER (yellow), regulation of cell death (blue),translation (pink),melanosome(brown),myelins heath (aqua),and muscle contraction (amber).

Eleven altered proteins were identified in all 6 patients, including known autoAgs (ACTB, EEF1A1, EEF2, ENO1, LGALS1, PABPC1) and unknown ones (CRTAP, NAP1L1, PSAP, RRBP1, TPT1) (Table 1). AHNAK (neuroblast differentiation-associated protein, a known autoAg in lupus) was identified in 5 patients. Overall, a majority of the altered proteins identified from the 6 COVID patients are known autoAgs, e.g., CALM1, CALR, CALU, CANX, DNAJB11, HDGF, HSPA5 (BiP), IQGAP1, LCP1, LMNB1, MYH9, NACA, P4HB, SFPQ, PDIA3, TPM3, TUBB, VCP, VIM, WARS, and YB3 (Table 1). Unknown or putative autoAgs include CAP1, CTSB, HDLBP, HYOU1, SND1, and SUB1.

We initially identified 30 DS-affinity proteins from HFL1 cells related to the melanosome, and, intriguingly, all of these are also COVID-altered proteins (Fig. 5B). Based on STRING GO analysis, the melanosome is the most significant cellular component related to altered proteins in all 6 patients (with false discovery rates ranging from 1.52E-8 to 1.11E-23). In HIV infection, melanosome production is stimulated in some patients and leads to an increase in pigmented lesions (*69*). However, melanosome involvement in COVID-19 is not known. Two Wuhan doctors in intensive care for COVID temporally turned dark, although the cause was thought to be a drug reaction. A COVID patient has been reported with acute flaccid tetraparesis and maculopapular pigmented plaques on the limbs (*70*). In mice, coronavirus induces an acute and long-lasting retinal disease, with initial retinal vasculitis followed by retinal degeneration that is associated with retinal autoantibodies and retinal pigment epithelium autoantibodies (*71*).

### Association between autoimmunity and virus infections

We identified COVID-altered proteins with DS-affinity that are involved in the host response to various aspects of viral infection and that possess a high propensity to become autoAgs. For example, viral RNA metabolism, translation, vesicles, and vesicle transport contribute a large number of known and putative autoAgs. In addition, viral processes, particularly symbiont processes and interspecies interactions between host and viruses, contribute significantly to altered proteins (Fig. 7A). For example, among altered proteins related to response to viral processes, HSPA8, DDB1, RAD23A, PABPC1, PPIB, P4HB, LGALS1, GSN, and ILF3 are known autoAgs (Table 1).

**Fig. 7A.**
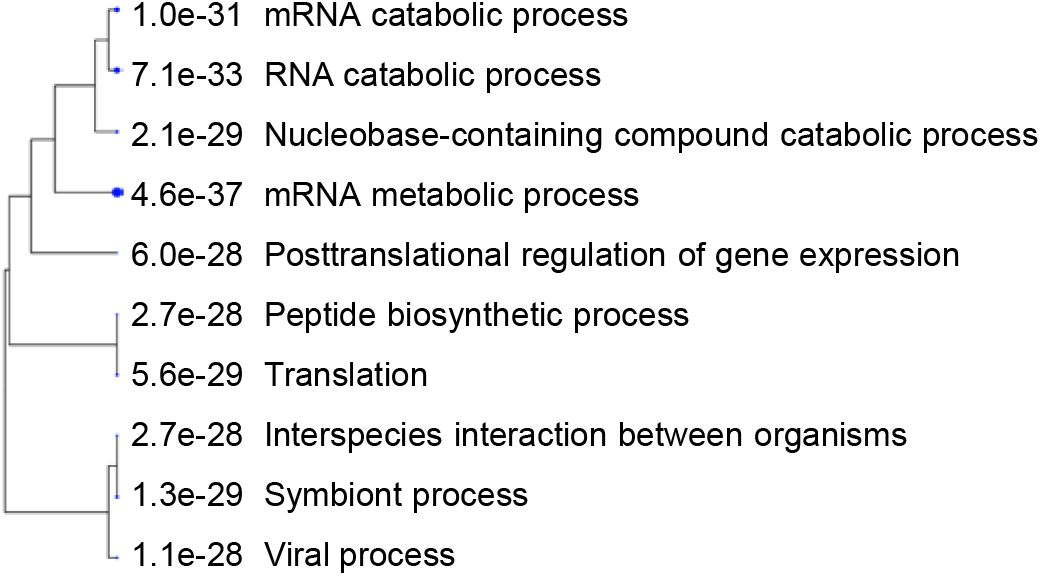
Hierarchical clustering of top 10 pathways involving COVID-altered proteins. Analysis based on hypergeometric distribution followed by FDR correction.

In particular, COVID-altered cytoskeletal filament proteins shed light on viral trafficking in host cells. SARS-CoV-2 infection induces profound remodeling of the cytoskeleton, and replicating viral vesicles are surrounded by a network of intermediate filaments (*72*). The cytoskeletal network appears to facilitate coronavirus transport and expulsion, with thickening actin filaments providing the bending force to extrude viral vesicles (*73*). We identified 84 altered proteins related to the cytoskeleton and 84 altered proteins related to vesicle-mediated transport (Fig. 2). These altered proteins are implicated in various processes, including cytoskeleton-dependent intracellular transport, actin fiber-based movement, actin-mediated cell contraction, microtubule-dependent trafficking from the Golgi to the plasma membrane, and transport along microtubules.

Many positive-strand RNA viruses (including SARS-CoV-2, Enterovirus, Hepatitis C virus, Norovirus, and Poliovirus) hijack a common group of nuclear factors to support the biosynthetic functions required for viral replication and propagation (*74*). 20 of these hijacked nuclear proteins are identified by DS-affinity in our study (Fig. 7). In addition, altered proteins are found in other viral infections, including porcine reproductive and respiratory syndrome virus (*75*), H5N1 avian influenza viruses (*76, 77*), Japanese encephalitis virus (*54*), Rift Valley fever virus (*78*), Hepatitis B virus (*79*), HIV (*80–82*), Herpes Simplex virus (*83*), and Epstein-Barr virus infection (Fig. 7B and STRING ontology analysis).

**Fig. 7B.**
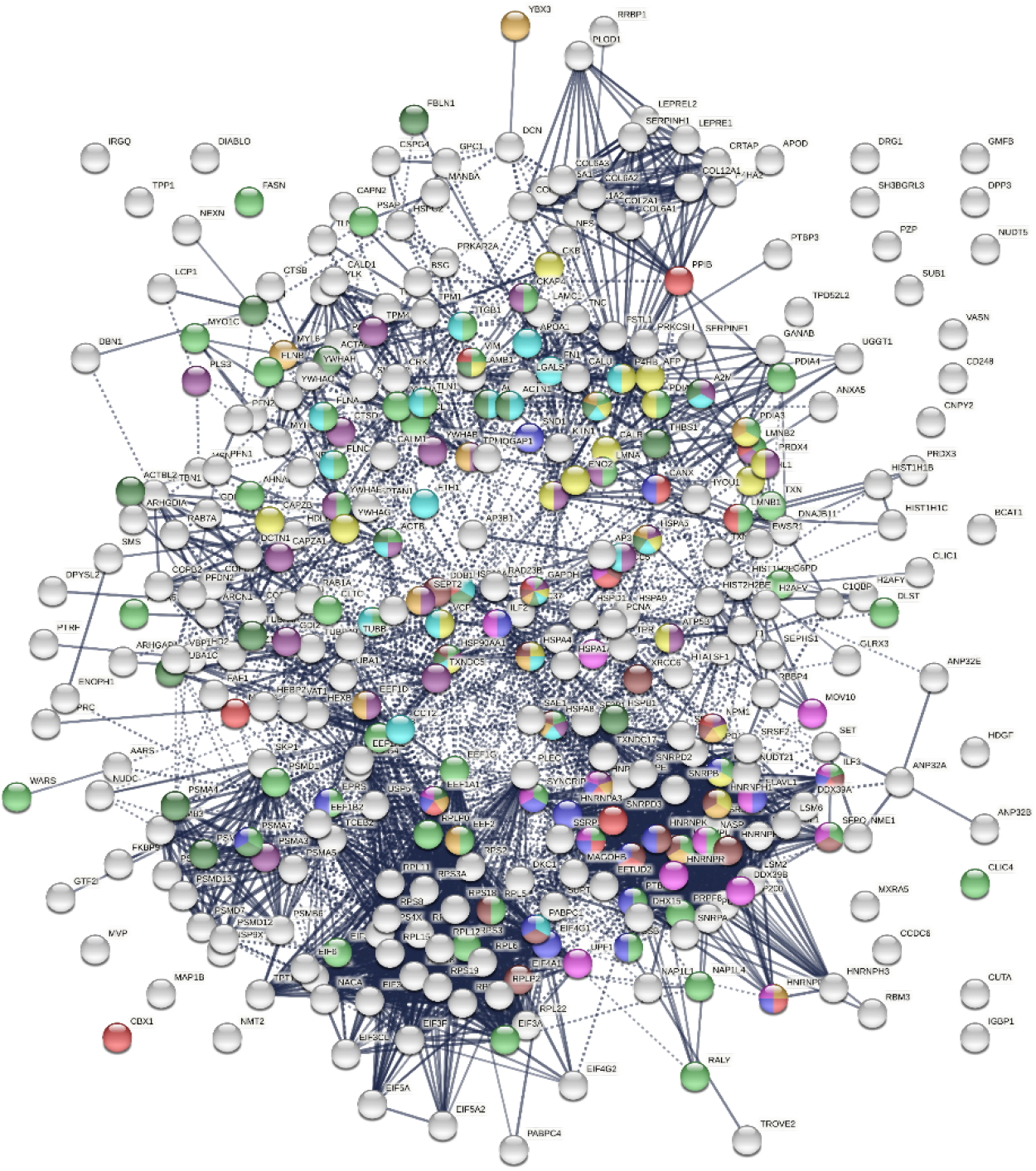
COVID-altered host proteins with DS-affinity found in various viral infections. Porcine reproductive and respiratory syndrome (56 proteins, green), H5N1 avian influenza virus (27 proteins, dark purple), Japanese encephalitis virus (23 proteins, gold), Rift Valley fever virus (24 proteins, aqua), Hepatitis B virus (22 proteins, dark green), HIV (identified in different studies, 18 amber, 18 brown, 18 red and 17 pink), and shared among positive-sense RNA viruses (20 proteins, blue).

### Autoimmunity concerns for mRNA vaccines

Our study identified a large number of known and putative autoAgs that are related to mRNA metabolism, translation, vesicles, and vesicle trafficking (Figs. 1–2). This finding begs us to wonder whether mRNA vaccines may induce unintended autoimmune consequences in the long term. mRNA vaccines are essentially synthetic viral vesicles. To induce protective immunity, mRNA vaccine vesicles will need to be transported into cells where they hijack the host cell machinery to produce a viral protein antigen, whereupon the antigen will be processed and presented by MHC molecules to induce B and T cell responses.

mRNA translation requires ribosomes, translation initiation factors, aminoacyl-tRNA synthetases, and elongation factors. We identified 18 ribosomal proteins by DS-affinity, all of which are altered in SARS-CoV-2 infection and 9 of which are known autoAgs (see references in Table 1). We also identified 15 eukaryotic translation initiation factor proteins, with 12 of them being COVID-altered and 4 being known autoAgs (Table 1). Six elongation factor proteins (5 subunits of EEF1 complex, EEF2) were identified by DS-affinity, of which all 6 are COVID-altered and 3 are known autoAgs (Table 1). Six tRNA synthetases were identified, with 5 being known autoAgs and 3 (AARS, EPRS, WARS) COVID-altered (Table 1). Autoantibodies to AARS are associated with interstitial lung disease and myositis (*84, 85*). EPRS appears to regulate pro-fibrotic protein synthesis during cardiac fibrosis (*86*). Gene mutations of WARS cause an autosomal dominant neurologic disorder characterized by slowly progressive distal muscle weakness and atrophy affecting both the lower and upper limbs (*87, 88*).

Once synthesized, the exogenous protein antigens are degraded by proteasomes, and the resulting peptides are transported into the ER where they are loaded onto MHC molecules by peptide loading complexes for presentation to T cells. In relation to these steps, 15 proteasome subunits were identified by DS-affinity, with 12 being COVID-altered and 7 being known autoAgs (Table 1). Nine proteins related to antigen processing and presentation are found to be altered in the 6 COVID-19 patients analyzed in this study, including HSPA1A, HSPA8, HSP90AA1, HSPAB1, HSPA5, PDIA3, CANX, CALR, and CTSB, with 7 being known autoAgs (Fig. 5 and Table 1).

In addition, among the 352 COVID-altered proteins identified in this study, 69 proteins are associated with mRNA metabolism (Fig. 2). Many of these proteins may be irrelevant to non-replicating mRNA molecules in mRNA vaccines, however, some are likely needed in processes such as 3’ end processing, deadenylation, and nonsense-mediated decay. For example, we identified poly(A) tail binding proteins PABPC1 and PABPC4 as COVID-altered proteins, both of which have been reported as autoAgs (Table 1).

mRNA vaccines are synthetic vesicles. This study identified 99 altered proteins associated with vesicles and 84 proteins associated with vesicle-mediated transport (Figs. 1, 2, 5). Although it is not clear which host molecules are involved in extra- and intracellular transport and uptake of mRNA vaccine vesicles, some of the vesicle-related proteins identified as DS-affinity proteins may be involved, e.g., proteins of receptor-mediated endocytosis (APOA1, CALR, CANX, CAP1, CLTC, HSP90AA1, HSP90B1, HSPG2, ITGB1, YWHAH) or phagocytosis (ACTB, CRK, GSN, HSP90AA1, HSP90AB1, MYH9, MYO1C, PDIA6, RAB7A, THBS1, TXNDC5).

Overall, a significant number of autoAgs related to different steps of mRNA vaccine action were identified in this study; however, our findings do not mean that these autoAgs will lead to aberrant autoimmune reactions as a result of mRNA vaccination. The development of autoimmune diseases or autoimmunity-related diseases entails a complex cascade of molecular and cellular interactions. Long-term monitoring of autoimmune adverse effects will be needed.

## Conclusion

This study identifies an autoantigenome of 408 proteins from human fetal lung fibroblast HFL1 cells by DS-affinity and protein sequencing, of which at least 231 proteins are confirmed autoAgs. Of these, 352 (86.3%) are found to be altered in SARS-CoV-2 infection when compared to published data, with at least 210 COVID-altered proteins being known autoAgs. The altered proteins are significantly enriched in a number of pathways and processes and are closely connected to various disease manifestations of COVID-19, particularly neurological problems, fibrosis, muscle dysfunction, and thrombosis.

Viral infections cause significant perturbations of normal cellular and tissue component molecules in the host, leading to cell death and tissue injury. Autoantigens resulting from molecular alterations may result directly from the injury or indirectly from responses to the injury. As a stress response, DS biosynthesis may be ramped up to facilitate wound healing and dead cell clearance. DS associates with autoAgs and stimulates autoreactive B cells and autoantibody production. Specific autoantibodies that are initially induced in response to a certain injury site may circulate and attack secondary sites where the autoAgs are also expressed, leading to a complex array of local and systemic autoimmune diseases.

This study illustrates a strong connection between viral infection and autoimmunity. The COVID-19 autoantigenome provides a detailed molecular map for investigating the diverse spectrum of autoimmune sequelae caused by the pandemic. The autoantigen atlas we are establishing may also serve as a detailed molecular reference for monitoring possible autoimmune reactions to mRNA vaccines and other viral infections.

## Materials and Methods

### HFL1 cell culture

The HFL1 cell line was obtained from the ATCC (Manassas, VA, USA) and cultured in Eagle’s Minimum Essential Medium supplemented with 10% fetal bovine serum (Thermo Fisher) and a penicillin-streptomycin-glutamine mixture (Thermo Fisher) at 37 °C.

### Protein extraction

About 100 million cells were harvested and suspended in 10 ml of 50 mM phosphate buffer (pH 7.4) containing the Roche Complete Mini protease inhibitor cocktail. Cells were homogenized on ice with a microprobe sonicator until the turbid mixture became nearly clear with no visible cells left. The homogenate was centrifuged at 10,000 g at 4 °C for 20 min, and the supernatant was collected as the total protein extract. Protein concentration was measured with the RC DC protein assay (Bio-Rad).

### DS-Sepharose resin preparation

20 ml of EAH Sepharose 4B resins (GE Healthcare Life Sciences) were washed with distilled water three times and mixed with 100 mg of DS (Sigma-Aldrich) in 10 ml of 0.1 M MES buffer, pH 5.0. 500 mg of N-(3-dimethylaminopropyl)-N’-ethylcarbodiimide hydrochloride (Sigma-Aldrich) powder was added to the mixture. The reaction proceeded by end-over-end rotation at 25 °C for 16 h. After coupling, resins were washed with water and equilibrated first with a low-pH buffer (0.1 M acetate, 0.5 M NaCl, pH 5.0) and then with a high-pH buffer (0.1 M Tris, 0.5 M NaCl, pH 8.0).

### DS-affinity fractionation

The total proteins extracted from HFL1 cells were fractionated on DS-Sepharose columns with a BioLogic Duo-Flow system (Bio-Rad). About 40 mg of proteins in 40 ml of 10 mM phosphate buffer (pH 7.4; buffer A) were loaded onto the column at a rate of 1 ml/min. Unbound proteins were washed off with 60 ml of buffer A, and weakly bound proteins were eluted with 40 ml of 0.2 M NaCl in buffer A. DS-binding proteins were eluted with sequential 40-ml step gradients of 0.5 M and 1.0 M NaCl in buffer A. Fractions were desalted and concentrated to 0.5 ml with 5-kDa cut-off Vivaspin centrifugal filters (Sartorius). Fractionated proteins were separated by 1-D SDS-PAGE in 4-12% Bis-Tris gels, and the gel lanes corresponding to 1.0 M or 0.5 M NaCl elutions were divided into two or three sections for sequencing.

### Mass spectrometry sequencing

Protein sequencing was performed at the Taplin Biological Mass Spectrometry Facility at Harvard Medical School. Proteins in gels were digested with sequencing-grade trypsin (Promega) at 4 °C for 45 min. Tryptic peptides were separated on a nano-scale C18 HPLC capillary column and analyzed in an LTQ linear ion-trap mass spectrometer (Thermo Fisher). Peptide sequences and protein identities were assigned by matching the measured fragmentation pattern with proteins or translated nucleotide databases using Sequest. All data were manually inspected. Only proteins with ≥2 peptide matches were considered positively identified.

### COVID data comparison with Coronascape

DS-affinity proteins were compared with currently available proteomic and transcriptomic data from SARS-CoV-2 infection compiled in the Coronascape database (as of 12/14/2020) (*28–48*). These data had been obtained with proteomics, phosphoproteomics, interactome, ubiquitome, and RNA-seq techniques. Up- and down-regulated proteins or genes were identified by comparing COVID-19 patients vs. healthy controls and cells infected vs. uninfected by SARS-CoV-2. Similarity searches were conducted between our data and the Coronascape database to identify DS-affinity proteins (or their corresponding genes) that are up- and/or down-regulated in the viral infection.

### Pathway and process enrichment analysis

Pathways and processes enriched in the putative autoantigenome were analyzed with Metascape (*28*). The analysis was performed with various ontology sources, including KEGG Pathway, GO Biological Process, Reactome Gene Sets, Canonical Pathways, CORUM, TRRUST, and DiGenBase. All genes in the genome were used as the enrichment background. Terms with a p-value <0.01, a minimum count of 3, and an enrichment factor (ratio between the observed counts and the counts expected by chance) >1.5 were collected and grouped into clusters based on their membership similarities. The most statistically significant term within a cluster was chosen to represent the cluster. Pathway hierarchical clustering was obtained with ShinyGo (*89*).

### Protein-protein interaction network analysis

Protein-protein interactions among collections of DS-affinity proteins were analyzed by STRING (*51*), including both direct physical interaction and indirect functional associations. Interactions are derived from genomic context predictions, high-throughput lab experiments, co-expression, automated text mining, and previous knowledge in databases. Each interaction is annotated with a confidence score from 0 to 1, with 1 being the highest, indicating the likelihood of an interaction to be true. Only interactions with high confidence (a minimum score of 0.7) are shown in the figures.

### Literature text mining

Literature searches in Pubmed were performed for every DS-affinity protein identified in this study. Search keywords included the protein name, its gene symbol, alternative names and symbols, and the MeSH keyword “autoantibodies”. Only proteins with their specific autoantibodies reported in PubMed-listed journal articles were considered “confirmed” autoAgs in this study.

## Acknowledgements and funding statement

This work was partially supported by Curandis, the US NIH, and a Cycle for Survival Innovation Grant (to MHR). MHR acknowledges the NIH/NCI R21 CA251992 and MSKCC Cancer Center Support Grant P30 CA008748. The funding bodies were not involved in the design of the study and the collection, analysis, and interpretation of data. We thank Jung-hyun Rho for technical assistance with experiments. We thank Ross Tomaino and the Taplin Biological Mass Spectrometry facility of Harvard Medical School for expert service with protein sequencing.

## Authors’ contributions

JYW directed the study, analyzed data, and wrote the manuscript. WZ performed some experiments and reviewed the manuscript. MWR and BVR assisted in data analysis and manuscript preparation. MHR consulted on the study and edited the manuscript. All authors have approved the manuscript.

